# Machine Learning Guided Video Analysis Identifies Sound-Evoked Pain Behaviors from Facial Grimace and Body Cues in Mice

**DOI:** 10.1101/2025.09.30.679579

**Authors:** Benjamin J. Seicol, Amelie Valles, Anna Kohler, Elisabeth Glowatzki, Megan B. Wood

## Abstract

Humans can experience sound-evoked pain, either from extremely loud sounds or in cases of pain hyperacusis from typically tolerable sounds. However, the mechanisms underlying sound-evoked pain remain poorly understood. Developing behavioral methods to measure sound-evoked pain in animal models is critical for elucidating these mechanisms. Here, a machine learning-based approach was developed to measure sound-evoked pain in freely moving mice by analyzing facial grimace and body position from video recordings during sound exposure. Facial grimace, a commonly used method to detect pain in mice, and body position, which can be used to measure postural and movement changes that also indicate pain, were both quantified using a deep neural network model trained to extract established facial and body features from video recorded by a single camera. To validate the model’s capability to detect pain, a known painful state, migraine induced by the injection of the neuropeptide calcitonin gene-related peptide (CGRP), was used. Using this machine learning-based approach, the ability to quantify a pain response from CGRP-induced migraine, distinct from baseline behavior, was demonstrated, resulting in a defined pain threshold. Sound exposures at high intensities elicited significant changes in facial grimace and body position, in comparison, surpassing the pain threshold calculated from CGRP-induced migraine. These behavioral changes were absent in *Tmie*-knockout mice, which lack functional sound transduction in the cochlea. This automated, high-throughput framework enables objective and sensitive analysis of pain providing a foundation for future studies investigating the peripheral and central mechanisms of sound-evoked pain.

**Significance:** This study introduces a quantitative framework for assessing pain using a single-camera setup and machine learning guided analysis to capture and analyze mouse behavior. By integrating two established pain metrics, facial grimace and attenuated movement, this method enables precise, non-invasive quantification of pain-related behaviors. The approach was validated with a well-characterized pain model, migraine, induced by injection of the neuropeptide CGRP, demonstrating the ability to quantify a pain response distinct from baseline behavior. By applying this framework to sound-evoked pain, the data revealed that exposure to intense sound triggers significant pain behavioral responses. These novel findings provide insights into the behavioral manifestations and neural underpinnings of sound-evoked pain, offering a robust tool for studying the mechanisms of pain perception.

## Introduction

Humans can experience pain in response to sound. Such pain can either occur as a reaction to loud sounds (greater than 120 dB SPL; 1-4) or, in the case of pain hyperacusis, can be a response to typically tolerable noise exposures (5–7). When sound exposure becomes painful, the mechanisms that lead to sound-evoked pain remain unknown (8,9). Processing of sound-evoked pain information, for example, may involve activation of brain regions canonically associated with pain, such as the thalamus, along with paradoxical hypoactivity in the auditory cortex (10–12). Developing sensitive behavioral methods to measure sound-evoked pain in laboratory animals will help to determine whether animal models recapitulate human pathology and allow further investigation of peripheral and central mechanisms responsible for sound-evoked pain (8, 13, 14).

Behavioral reflexes in response to sound onset are widely used to assess animals’ auditory sensitivity, for example by the acoustic startle response (4, 15, 16). Broadly, pain can be measured in mice either by 1) withdrawal reflex, such as, to thermal or mechanical stimuli (17), 2) operant conditions of forced-choice tasks (18), or 3) assessment of behavioral signatures associated with affective pain (e.g., facial grimace; 19, movement, 20). During pain perception, mice ‘grimace’, like humans (21, 22), and therefore pain can be quantified by characteristic changes in their facial features, for example, by eyes squinting (23). Facial grimace has effectively been used to measure painful states in animal models, for example in the case of migraine (24). Automated facial grimace detection and body movement analysis of persistent pain in mice, including post-operative and migraine pain, has proven comparable to individual investigators scoring behavioral cues; it has reduced bias, and allowed for high-throughput analysis of large datasets (20, 25, 26). Body movement has also been analyzed parallel, however, measuring behavioral changes in facial grimace and body position has required two or more camera angles (en face or profile and from above) (20, 27).

Due to the distinct behavioral approaches used to measure pain in animals, different types of pain are often defined as either ‘reflexive’ or ‘spontaneous’, where reflexive pain usually refers to a withdrawal reflex to an acute external sensory stimulus. In contrast, spontaneous pain is typically used to describe ongoing painful states (28). Pain evoked by sound exposures that persists for periods of time on the order of minutes is, therefore, a type of spontaneous pain (herein now referred to as “persistent pain”). Here, a deep neural network was trained to extract formerly established facial features (23) from video recordings of freely moving mice exposed to various levels of sound intensity presentations. Secondly, body position measurements from the same camera angle as facial grimace recordings were added to the analysis to simultaneously detect changes in mouse posture and locomotion behaviors associated with pain (29). This approach of analyzing combined facial grimace and body posture was validated with a well-characterized pain model, migraine, induced by injection of the neuropeptide CGRP, which demonstrated the ability to quantify a pain response distinct from baseline behavior and define a pain threshold. Sound exposure at high intensities elicited significant changes in facial grimace and body posture and, in comparison, surpassed the pain threshold established during migraine validation. Such changes did not occur in *Tmie*-knockout (*Tmie^-/-^)* mice that lack normal cochlear transduction of sound (30–33) validating that normal cochlear transduction is needed to induce sound-evoked pain behavior in mice. Facial grimace measurements combined with body position information extracted by deep learning from video recordings of mice presented with sound allowed for sensitive, objective, and high-throughput analysis of mouse sound-evoked pain.

## Results

Detecting persistent pain in mice has been performed before by analyzing facial grimace (21–23, 25, 26). Here, facial grimace detection has been adapted from work that has used six defined features of the face from the profile of a mouse’s face from videos of freely moving mice (23). It was further possible to expand beyond facial grimace using the same videos and some of these same markers to analyze body position information of the mice. Average values of each facial grimace parameter and separately of four specific body position parameters were calculated within defined periods (epochs) and combined to create an inventory of pain. This inventory was validated, and pain thresholds were determined, based on a known painful stimulus (CGRP-induced migraine). This approach was subsequently used to test the level of sound that elicits pain in mice. Additionally, mice without normal cochlear function were used to determine whether sound transduction in the inner ear is necessary for the generation of sound-evoked pain.

### Detection of facial grimace parameters in freely moving mice using a deep neural network

A deep neural network was trained to measure changes in facial grimace by automatic placement of thirteen points on the face of mice with skeletons connecting these points from video recordings of mice performed in controlled conditions (Figure 1). This approach allowed for high-throughput analysis of thousands of video frames from hours of recordings, greatly reducing the analysis time compared to manual annotation by blind observers as done before with the mouse grimace scale (21). In this study, facial grimace measurements were taken from mice enclosed in a custom-built clear chamber with scent holes on only one side that encouraged mice to remain mostly in profile with respect to the camera (Figure 1A). Representative images of mice in pain vs. control conditions illustrate the changes in the face, for example, facial grimace of a mouse in a pain state manifests with clear changes in ear positions and eye squinting, as well as subtle changes in snout and mouth positions (Figure 1B). Changes can also be seen in body position, such as the mouse being hunched over and the head angle pointing down, which will be described in detail below. Based on the video recordings, six facial parameters (23) were calculated from the pose estimated points on the mouse’s face (Figure 1C).

**Figure 1.**
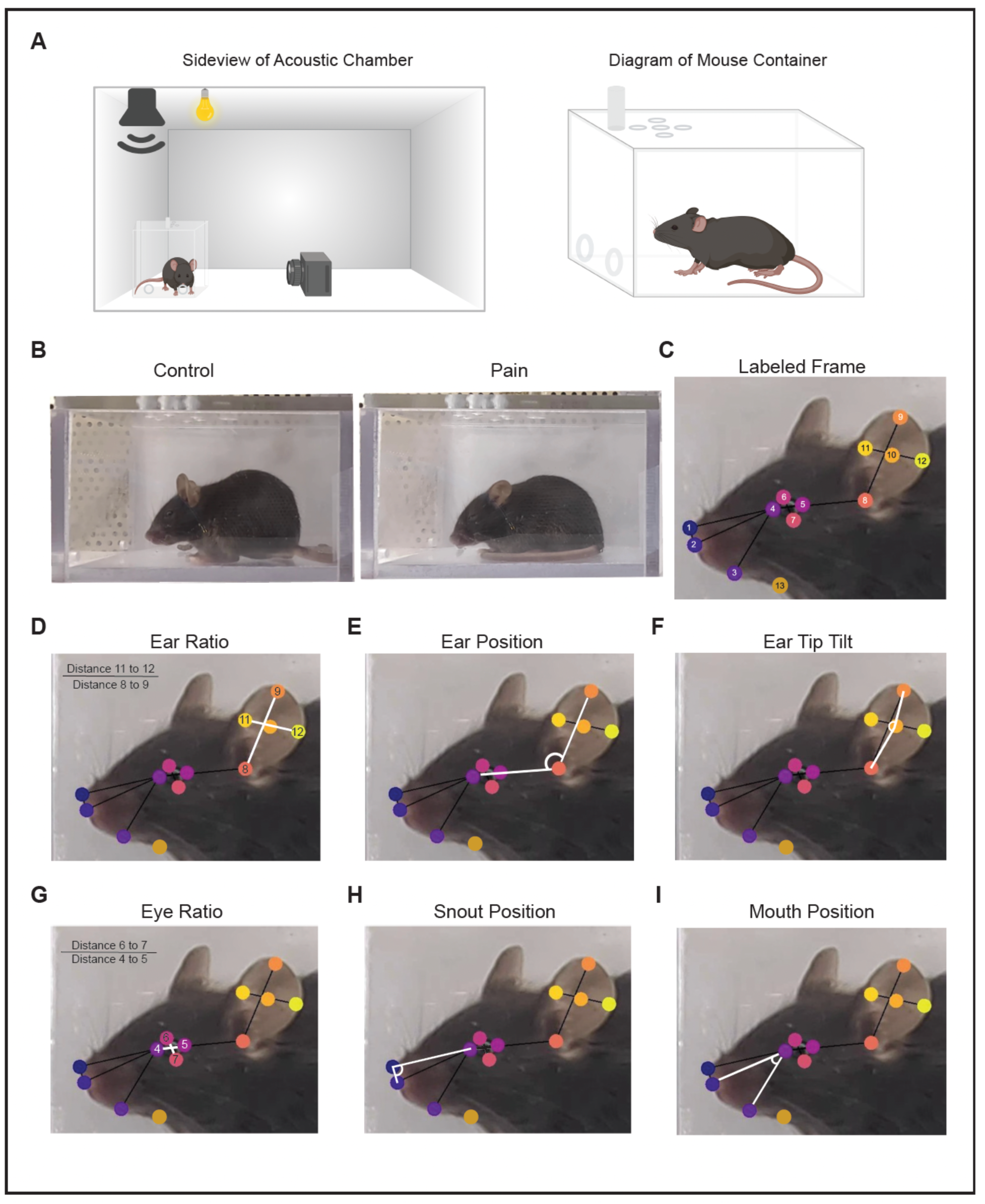
Detection of facial grimace parameters in freely moving mice using a deep neural network. **A**) Left, a schematic of the inside of the acoustic booth used for facial grimace video recording. Right, a diagram of the custom-built enclosure from the view of the camera. The ovals on the left wall and top indicate scent holes. Figure not to scale. **B**) Labeled frames from the same video of a mouse in a control condition (left) and a painful condition (right). **C**) A labeled frame of a mouse face showing the 13 markers on the face (numbered, colored dots). The black lines are skeletons generated by the model. Colored dot 1- Nose Tip (upper tip of nose); 2 – Nose Bot (bottom of nose); 3 – mouth; 4 – inner eye (anterior edge of eye); 5 – Outer Eye (caudal edge of eye); 6 – Eye Up (highest point of eye lid on dorsal edge); 7 – Eye Down (lowest point of eyelid on ventral edge); 8 – Ear canal (edge of hair and ear canal); 9 – Ear Tip (edge of pinna furthest from the ear canal); 10 – Ear center (center of ear pinna between points 9, 10, 11 and 12); 11 – Ear Med (halfway up the pinna on the medial side); 12 – Ear Lat (halfway up the pinna on the lateral side of the ear); 13 – Neck (point where the head connects to the body). **D**) Ear Ratio was calculated as the ratio of the length of the lines in pixels from Ear Med to Ear Lat (Point 11 to 12) and Ear Canal to Ear Tip (Point 8 to 9) indicated by white lines. **E**) Ear Position was calculated as the obtuse angle formed by lines connecting Inner Eye, Ear Canal, and Ear Tip with Ear Canal as the vertex as indicated in white. **F**) Ear Tip Tilt was calculated as the angle formed by lines connecting Ear Canal to Ear Center to Ear Tip with Ear Center as the vertex as indicated in white. **G**) Eye Ratio was calculated as the ratio of the length of the lines in pixels from Eye Up to Eye Down (Point 6 to7) and Inner Eye to Outer Eye indicated by white lines (Point 4 to 5). **H**) Snout Position was calculated as the acute angle formed by lines connecting Nose Tip, Inner Eye, and Nose Bottom with Inner Eye as the vertex as indicated in white. **I**) Mouth Position was calculated as the acute angle formed by lines connecting Nose Bottom, Inner Eye, and Mouth with Inner Eye as the vertex as indicated in white.

DeepLabCut software (DLC; (34, 35)) was used to train a neural network model capable of automatically identifying thirteen markers on the mice with skeletons connecting these points to calculate six facial parameters (Figure 1D-I). The changes in ear positions were mapped using 3 parameters: 1) ‘Ear Ratio’ (which goes up as the mouse’s ear lies flat against the head thereby increasing the width in relation to the camera in profile view; Figure 1D), 2) ‘Ear Position’ (which measures the angle that increases when the mouse’s ear rotates further towards its posterior; Figure 1E), and 3) ‘Ear Tip Tilt’ (which measures the angle of the top half of the ear relative to the bottom half which increases when the ear is flattened against the back of the head; Figure 1F). These three ways of measuring changes in ear position are valuable due to the manner of grimacing in animals that are exaggerated in the ears vs. humans (36–38). The three ear-related measures also each reflect a distinct neuromuscular circuit, arising from the 1) transverse auricular muscle, 2) interscutular muscle, and 3) anterior and posterior auricular levators, respectively (39, 40). Additional face grimace parameters measured were ‘Eye Ratio’ (which gets smaller with ‘eye squinting’; Figure 1G), ‘Snout Position’ (which gets smaller when the angle between the eyes, nose, and mouth is more acute meaning snout is more pointed during pain; Figure 1H) and Mouth Position’ (which gets smaller when the bottom lip extends outwards similar to clenching of the jaw; Figure 1I).

### Facial grimace scores in mice that experience migraine after CGRP injection indicate pain

Migraine can be elicited by injection of calcitonin gene-related peptide (CGRP; (41) and is a well-known painful stimulus in mice (42). Pain in this model has previously been detected using the mouse grimace scale, and peaks at approximately 30 minutes following CGRP administration (24). We therefore adopted this model to test our approach of detecting persistent pain with the set of facial grimace parameters described in Figure 1. Mice were injected with CGRP solution (100 µg/kg in saline, i.p.) or injected with saline (vehicle). Video recording began when mice were placed into the chamber for 10 minutes to record baseline behavior (‘*Baseline*’; t = −10 – 0 mins; Figure 2A). Mice then received an injection of either CGRP, or saline (t = 0 min), and the video recording was paused. Video acquisition was restarted 20 minutes after injection and continued for another 20 minutes (‘*All*’; t = 20 – 40 mins) to ensure complete capture of the expected migraine peak (Supplemental Video 1). Each facial grimace parameter measured during *Baseline* and *All* can be visualized as a diary plot of all analyzed frames at 30 frames/s, here shown for data from an individual mouse (e.g., Ear Position; Figure 2B). It is notable that there are gaps in the dairy plot illustrating periods where the points could not all be applied to the face, for example when the mouse was turned away from the camera (Supplemental Video 1). Data from all mice (N = 12) were superimposed and are shown in Figure 2C. The expected peak of migraine-like pain from CGRP was anticipated to be around t = 30 minutes. An empirical peak emerged from combined data of facial grimace and body position parameters (quantified below in Figure 5), showing that during a 5-minute epoch (‘*Peak*’; t = 27 – 32 minutes) surrounding the expected peak mice exhibited significant changes in behavior. This empirical peak was therefore analyzed separately from the remaining 15 minutes of CGRP recording (‘*Non-Peak*’), including the time before and after the ‘*Peak*’.

**Figure 2.**
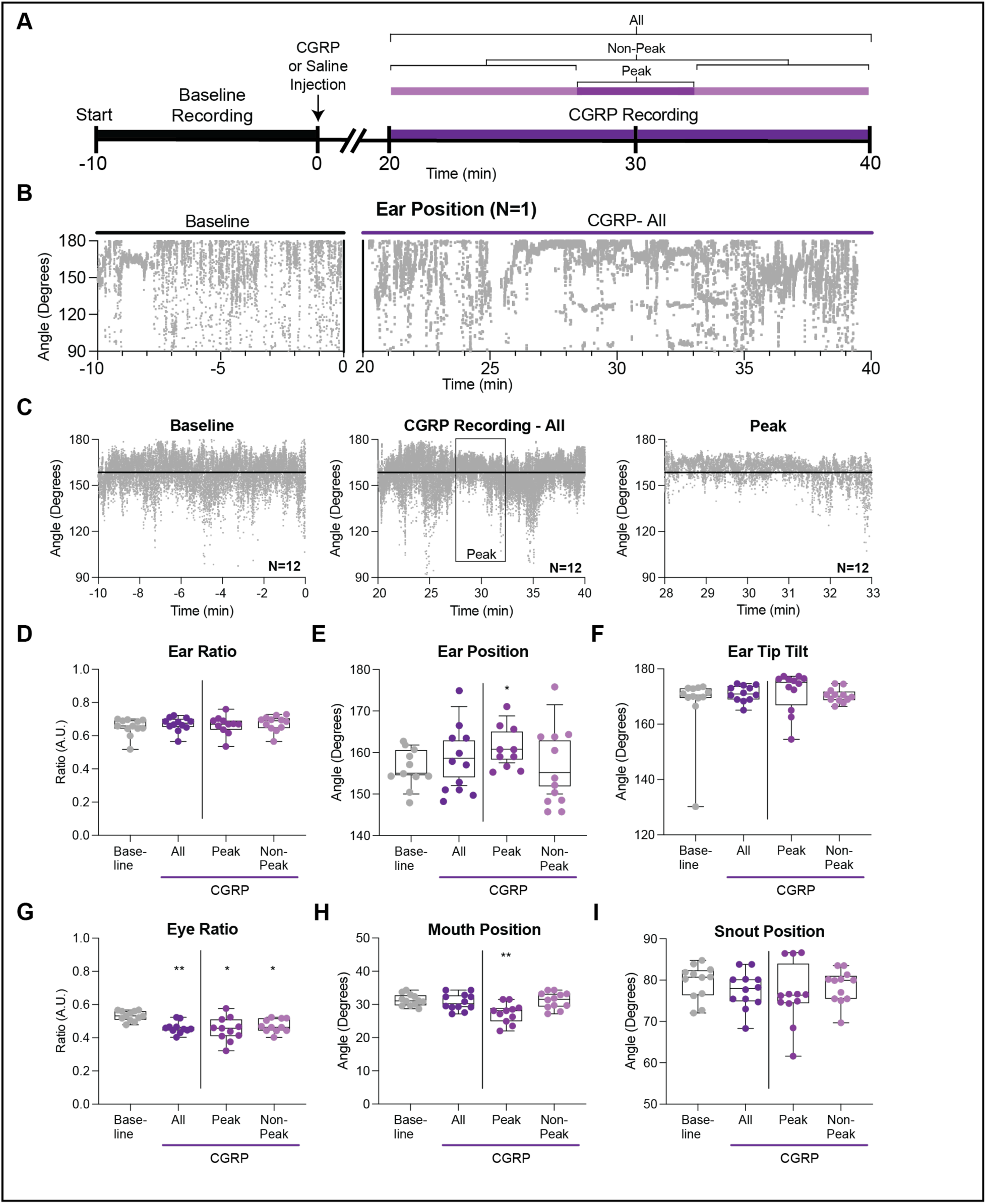
Facial grimace scores in mice that experience migraine after CGRP injection. **A**) Timeline of CGRP injection experiment. Animals were filmed for a 10-minute baseline period. CGRP was injected based on body weight. After a 10-minute period, mice were filmed for 20 minutes surrounding the peak of action CGRP which occurs 30 minutes after injection. **B**) Example of the Ear Position data from the baseline and CGRP time periods of one mouse (N=1). Each dot is the measurement from a single frame of video. **C**) Averaged data for each frame of video for Ear Position from 12 mice (N=12) during Baseline and the entire CGRP Recording. The right graph shows the average Ear Position for 12 mice during the peak time period as indicated in A and the box in the middle graph. The black horizontal line is the average of the baseline time-period as a reference in all graphs. (**D-I**) The following graphs show the averaged data for each facial parameter indicated. Each dot is the averaged data for the indicated time-period for 1 mouse. *Baseline* is the averaged data for each mouse for all frames from the Baseline video (10 minutes). ‘*All*’ is the averaged data for each mouse for all frames after CGRP injection. ‘*Peak*’ is the averaged data from minutes 27-31 and after CGRP injection. ‘*Non-Peak*’ is the averaged data from time outside of the peak time period (minutes 20-26 and minutes 32 to 40). Box and whiskers have a line at the median and the box is the interquartile range with the whiskers extending to the maximum and minimum value of the data. **D**) Ear Ratio. **E**) Ear Position. **F**) Ear Tip Tilt. **G**) Eye Ratio, *Baseline* compared to *All*: paired, two-tailed t-test, **, p=0.0012; repeated measures mixed effects model, **, p=0.0025; post hoc comparisons using Tukey’s correction, *Baseline* compared to *Peak*, **, p= 0.0215; *Baseline* compared to *Non-Peak*, *, p= 0.0111. **H**) Mouth Position, repeated measures mixed effects model, ****, p=<0.0001; post hoc comparisons using Tukey’s correction, *Baseline* compared to *Peak*, **, p=0.0015; *Peak* compared to *Non-peak*, p=0.0026. **I**) Snout Position. If no statistics noted, data was not significantly different.

Median values from *Baseline* and *All* epochs were compared using paired, two-tailed, student t-test for each of the six facial grimace measurements, while *Baseline*, *Peak*, and *Non-Peak* epochs were compared using either repeated measures one-way ANOVA for balanced groups or repeated measures mixed effects model for unbalanced groups (Figure 2D-I). Significant differences, relative to *Baseline*, were found during *Peak* for Eye Ratio and Mouth Position. Eye Ratio also differed from *Baseline* during *All* and *Non-Peak*. Specifically, Eye Ratio was significantly reduced in *All* compared to *Baseline* (Difference, −0.0781 AU, paired t-test, p < 0.005). Eye Ratio was also significantly lower than *Baseline* in both *Peak* and *Non-Peak* epochs (*Baseline*, *Peak*, and *Non-Peak*, repeated measures mixed effects model, p = 0.0025; post hoc Tukey’s correction, *Peak* vs. *Baseline* median difference = −0.0742 AU, p < 0.05, and *Non-Peak* vs. *Baseline* median difference = −0.0681 AU, p < 0.05). Mouth Position was also significantly lower during *Peak* vs. *Baseline*, as well as during *Peak* vs. *Non-Peak* (repeated measures mixed effects model, p < 0.0001; post hoc comparisons using Tukey’s correction, *Peak* vs. *Baseline* median difference = −2.98°, p < 0.005, *Peak* vs. *Non-Peak* median difference = −3.49°, p < 0.005). No significant differences from *Baseline* or between epochs were found in Ear Ratio, Ear Position, Ear Tip Tilt, or Snout Position. Additionally, no significant differences were observed in any measures in saline-injected mice (Supplemental Figure 1).

### Body position parameters in mice experiencing migraine after CGRP injection

A subset of the pose estimated points used for measuring facial grimace were additionally applied to generate measurements of mouse body position that could indicate pain. Body position parameters were chosen based on the observation of changed locomotive behaviors (43), whereby mice were observed to explore, rear, and generally move less during pain states, as well as maintain a hunched posture more often. Body position measurements were therefore quantified to grossly incorporate changes in these complex behaviors (Figure 3), namely, the ‘Relative Nose Tip Position’ (Figure 3A; related to exploring), ‘Percentage of Nose Tip in Top 1/3 of Chamber’ (Figure 3B; captured rearing), ‘Vertical Line Crosses of Nose Tip’ (Figure 3C; measured turning), and ‘Face Inclination as Slope of Nose Tip to Neck’ (Figure 3D; showed hunching).

**Figure 3.**
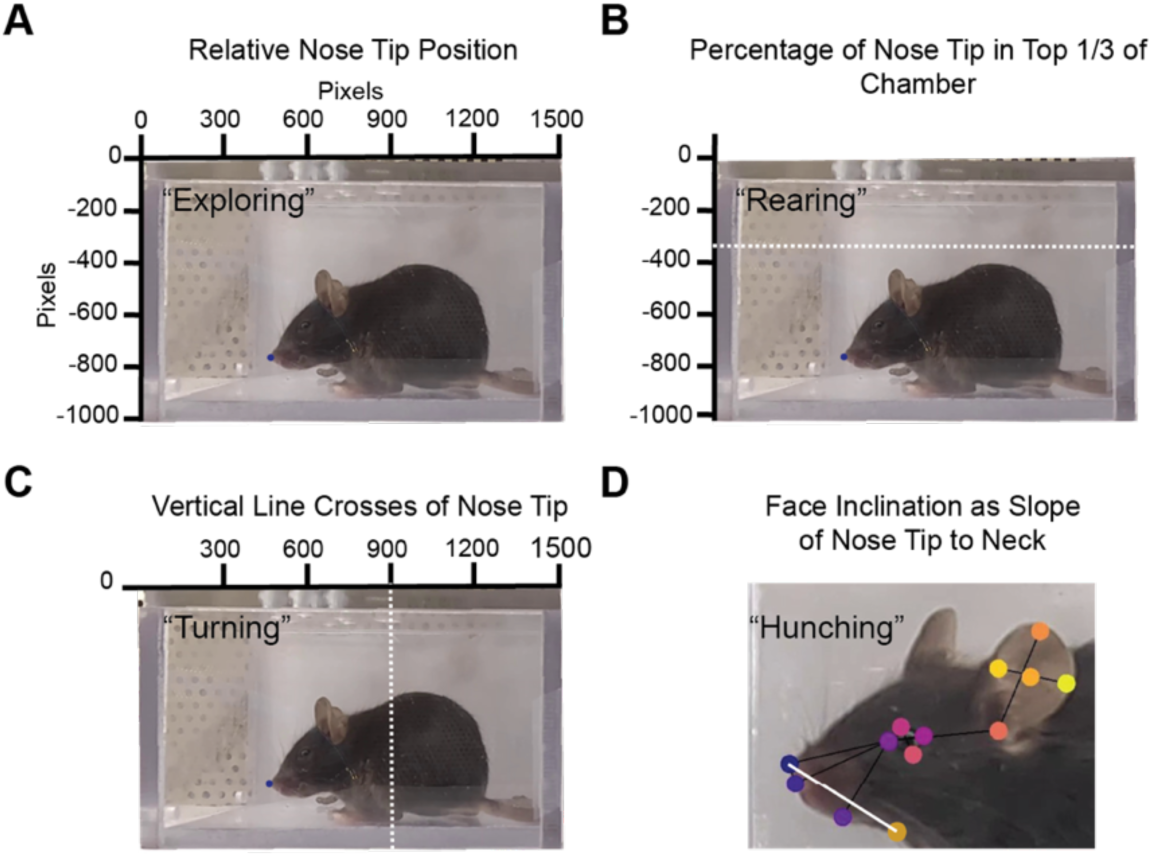
Detection of body position parameters in freely moving mice using a deep neural network. **A**) Example of a full frame of video of a control mouse with x and y axes showing pixel values. The nose tip is indicated with a navy-blue dot. The y-axis pixel value of the nose tip was recorded for each frame and averaged over discrete time periods to create the measurement named Relative Nose Tip Position as a proxy for exploring behavior. **B**) Example of a full frame of video of a control mouse. The white dashed line at −450 pixels on the y-axis indicates the boundary of the top 1/3 of the chamber. The average number of frames in a time-period where the nose tip (navy-blue dot) was above this line based in the y-axis was calculated as the Nose Tip Top % measurement as a proxy for rearing behavior. **C**) Example of a full frame of video of a control mouse. The white dashed line at 900 pixels on the x-axis indicates a boundary which was used to count the number of times the nose tip (navy-blue dot) crossed this line in a time-period. This was used to calculate the Vertical Line Crosses measurement as a proxy for turning behavior. The line is intentionally offset to capture the full turning of the mouse from one side of the cage to the other instead of rearing straight backwards as during grooming. **D**) Example of a mouse face labeled from the DNN training. The y-axis value of the Nose Tip was subtracted from the y-axis value of the Neck point (Neck – Nose Tip; indicated by the white line). The Face Inclination measurement was then calculated as the percentage of frames during a time-period that the Neck – Nose Tip value was negative meaning the Nose Tip was below the Neck value on the Y-axis as a proxy for hunched body position.

Body position parameters (Figure 4) were plotted in the same way as facial grimace measurements. A representative diary plot shows relative nose tip position during migraine, (Figure 4A). Superimposed plots from all mice for Relative Nose Tip Position (shown in Figure 4B) illustrate a subtle decrease in variability during *Peak*. Appropriate statistical tests were performed to compare groups, and no significant differences were found for Relative Nose Tip Position (Figure 4C). In contrast, *Peak* was significantly different compared to *Baseline* for Percentage in Top (Figure 4D; repeated measures one-way ANOVA, p = 0.022; post hoc comparisons using Tukey’s correction, *Peak* vs. *Baseline* median difference = −8.25 percentage points, p < 0.05), for Vertical Line Crosses (Figure 4E, repeated measures mixed effects model, p = 0.0004; post hoc comparisons using Tukey’s correction, *Peak* vs. *Baseline* median difference = −39 incidence, p < 0.005), and for Face Inclination (Figure 4F, repeated measures mixed effects model, p = 0.03; post hoc comparisons using Tukey’s correction, *Peak* vs. *Baseline* median difference = 35.14 percentage points, p < 0.05). Reductions in *All*, compared to *Baseline* were found for both Vertical Line Crosses (paired t-test, *All* vs. *Baseline* median difference = −24.5 incidences, p < 0.05) and Face Inclination (paired t-test, *All* vs. *Baseline* median difference = 15.46 percentage points, p < 0.05). Additionally, Vertical Line Crosses was significantly decreased during *Peak* vs. *Non-Peak* (paired t-test, *Peak* vs. *Non-Peak* median difference = −19 incidences, p < 0.001).

**Figure 4.**
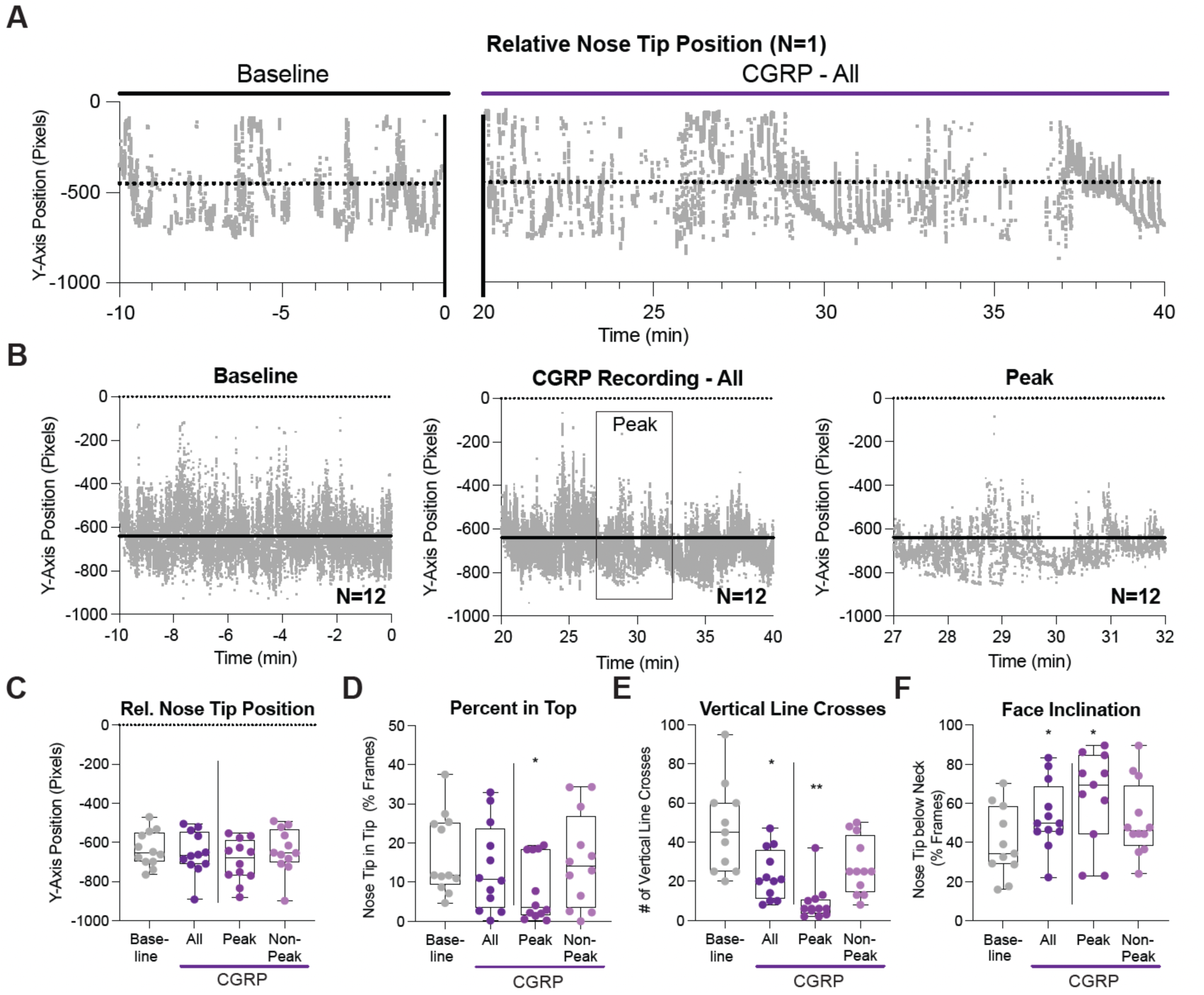
Body position parameters in mice experiencing migraine after CGRP injection. **A**) Example of the Relative Nose Tip Position data from the baseline and CGRP time periods of one mouse (N=1). Each dot is the measurement of Y-axis Nose Tip Position from a single frame of video. The line at Y-axis −450 was used to calculate the Percent in Top measurement. **B**) Averaged data for each frame of video for Relative Nose Tip Position from 12 mice (N=12) during Baseline and the entire CGRP Recording. The right graph shows the average Relative Nose Tip Position for 12 mice during the peak time period as indicated in Fig 2A and the box in the middle graph. The black horizontal line is the average of the baseline time-period as a reference in all graphs. **C-F**) The following graphs show the averaged data for each body position measurement indicated. Each dot is the averaged data for 1 mouse. *Baseline* is the averaged data for each mouse for all frames from the Baseline video. ‘*All*’ is the averaged data for each mouse for all frames after CGRP injection. ‘*Peak*’ is the averaged data from minutes 27-31 and after CGRP injection. ‘*Non-Peak*’ is the averaged data from time outside of the peak time period (minutes 20-26 and minutes 32 to 40). Box and whiskers have a line at the median and the box is the interquartile range with the whiskers extending to the maximum and minimum value of the data. **C**) Rel. Nose Tip Position. **D**) Percent in Top: *Baseline* compared to *Peak* and *Non-Peak*, repeated measures one-way ANOVA, p=0.0221; post hoc comparisons using Tukey’s correction, *Baseline* compared to *Non-Peak*, *, p= 0.0266. **E**) Vertical Line Crosses: The absolute number of crosses was normalized to a 5-minute time-period to compare baseline (10 min) and All (20 min) to Peak (5 min) and *Non-Peak* (15 min). *Baseline* compared to *All*: paired, two-tailed t-test, *, p=0.0241; *Baseline* compared to *Peak* and *Non-Peak*, repeated measures mixed effects model, **, p=0.0004; post hoc comparisons using Tukey’s correction, *Baseline* compared to *Peak*, ***, 0.0025; post hoc comparisons using Tukey’s correction, *Peak* compared to *Non-Peak*, ***, 0.0002. **F**) Face Inclination: *Baseline* compared to *All*: paired, two-tailed t-test, *, p=0.0236; *Baseline* compared to *Peak* and *Non-Peak*, repeated measures mixed effects model, *, p=0.03; post hoc comparisons using Tukey’s correction, *Baseline* compared to *Peak*, *, p=0.0293. If no statistics noted, data was not significantly different.

### Combined Facial Grimace and Body Position Scores provide detection of pain in mice

Individual facial grimace and body position parameters were combined to create two separate composite scores. Each measurement was first converted to a percentage change from *Baseline* (Supplemental Figure 2), and these were then added together to calculate the summed values for facial grimace or for body position for both CGRP- and saline-injected mice (Figure 5). CGRP-injected mice showed significant changes between *Baseline* and *All* (referring to all frames recorded) in the summed measurements for both Facial Grimace and Body Positions Scores, while saline-injected mice showed no changes (Figure 5A-B). Specifically, CGRP significantly lowered both Facial Grimace Score and Body Position Score in *All* relative to *Baseline* (Facial Grimace Score: paired t-tests, *All* vs. *Baseline* median difference = −20.59 percentage points, p < 0.005; Body Position Score: paired t-tests, *All* vs. *Baseline* median difference = −162.6 percentage points, p < 0.001). These data show that pain can be detected in both the composite scores of facial grimace and body position using this approach.

**Figure 5.**
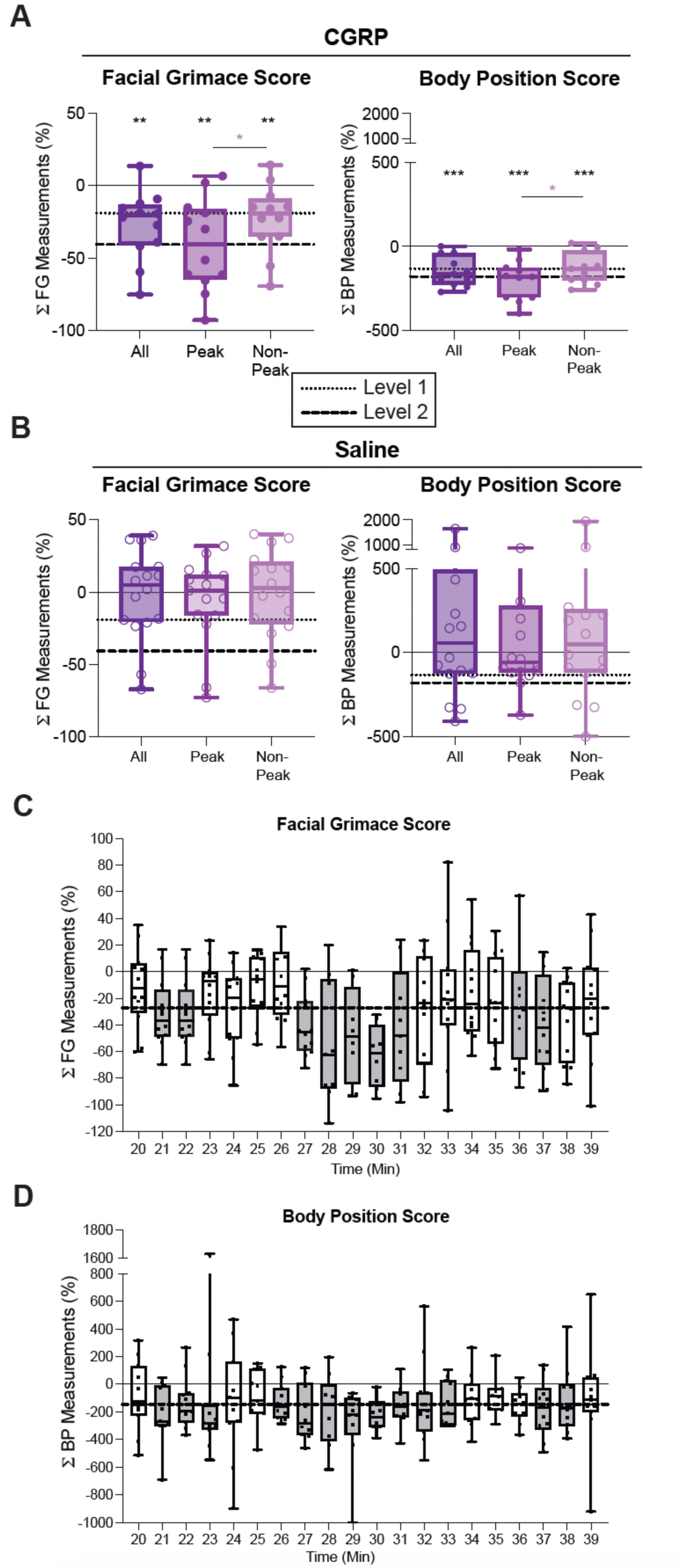
Combined Facial Grimace and Body Position Scores provide quantification of pain in mice. Facial Grimace Scores were calculated as the added percentage change from baseline for all facial grimace measurements for each mouse. When a facial grimace parameter resulted in a positive change from baseline (Ear Ratio, Ear Position), it was inverted so the percentage change from baseline associated with pain was a negative value. Body Position Scores were calculated as the added percentage change from baseline for all position measurements for each mouse. When a body position measurement resulted in a positive change from baseline (Rel. Nose Tip Position, Face Inclination), it was inverted so the percentage change from baseline associated with pain was a negative value as with the Facial Grimace Score. **A**) Facial Grimace (left) and Body Position (Right) Scores for mice injected with CGRP (N=12). Box and whiskers have a line at the median and the box is the interquartile range with the whiskers extending to the maximum and minimum value of the data. CGRP Facial Grimace Score: *Baseline* compared to *All*, paired, two-tailed t-test, **, p=0.0022; *Baseline* compared to *Peak*, paired, two-tailed t-test, **, p=0.001; *Baseline* compared to *Non-Peak*, paired, two-tailed t-test, *, p=0.0063. *Peak* compared to *Non-Peak*, paired, two-tailed, t-test, *., p=0.0263. The dotted line is defined as the score associated with Pain Level 1 and begins at the median of the *Non-Peak* data (median = −19.04 percent change from baseline). The dashed line is defined as the score associated with Pain Level 2 and begins at the mean of the *Peak* data (median = −40.62 percent change from baseline). CGRP Body Position Score (Right); Baseline compared to All, paired, two-tailed t-test,***, p=0.0003; *Baseline* compared to *Peak*, paired, two-tailed t-test, ***, p=0.002; Baseline compared to *Non-Peak*, paired, two-tailed t-test, *, p=0.001. *Peak* compared to *Non-Peak*, paired, two-tailed t-test,**, p=0.0369. Box and whiskers have a line at the median and the box is the interquartile range with the whiskers extending to the maximum and minimum value of the data. The dotted line is defined as the score associated with Pain Level 1 and begins at the median of the *Non-Peak* data (median = −132.4 percent change from baseline). The dashed line is defined as the score associated with Pain Level 2 and begins at the mean of the Peak data (median = −179.5 percent change from baseline). For a mouse to be experiencing pain, the scores must be lower than −19.04 for Facial Grimace and −132.4 for Position. **B**) Facial Grimace (Left) and Body Position (Right) Scores for mice injected with Saline (N=16). Box and whiskers have a line at the median and the box is the interquartile range with the whiskers extending to the maximum and minimum value of the data. Statistical tests were not significant. Saline Facial Grimace Score: Baseline compared to All, paired, two-tailed t-test, p=0.8211. Baseline compared to *Peak* and *Non-Peak*, repeated measures mixed model, p=0.5096. Saline Body Position Score: *Baseline* compared to *All*, paired, two-tailed t-test, p=0.1673. *Baseline* compared to *Peak* and *Non-Peak*, repeated measures mixed model, p=0.2735. **C**) The percentage change from baseline for each facial grimace measurement was added together and averaged for each minute of the CGRP recording. Each dot corresponds to 1 animal. The dashed line is the mean of all 20 minutes for all mice (−27.17 % change from baseline). Minutes where the median falls below the dashed line are colored gray. Minutes 27-31 show the longest string of median scores below the dashed line. **D**) The percentage change from baseline for each body position measurement was added together and averaged for each minute of the CGRP recording. Each dot corresponds to 1 animal. The dashed line is the median of all 20 minutes for all mice (−145.7 % change from baseline). Minutes where the median falls below the dashed line are colored gray. Minutes 27-34 have the longest string of lowest median scores.

### Defining two pain levels based on thresholds calculated during distinct time windows after inducing migraine

Migraine induced by the injection of CGRP has been reported to cause a peak in the pain intensity in mice at approximately 30 minutes post injection (24). While prior studies have demonstrated this peak using a lower sampling rate (e.g., 4 images per time point, acquired 20 seconds apart), the current approach uses a higher sampling rate and therefore provides a more detailed assessment of the timeline of behavioral changes after inducing migraine in mice. Using this new approach, the empirical *Peak* of migraine pain emerged around the expected peak from t = 27 – 32 minutes post injection of CGRP, further defined below (Figure 5C-D). The remaining time, both before and after the defined *Peak* time period, was combined to calculate the *Non-Peak* period. The composite scores were compared with *Baseline* for both *Peak* and *Non-Peak* periods (Figure 5A-B), where both *Peak* and *Non-Peak* differed significantly from *Baseline* (*Peak* vs. *Baseline* median difference = −40.62 percentage points, p = 0.001, and *Non-Peak* vs. *Baseline* median difference = −19.04 percentage points, p < 0.01; Body Position Score: *Peak* vs. *Baseline* median difference = − 179.5 percentage points, p < 0.001, and *Non-Peak* vs. *Baseline* median difference = −132.4 percentage points, p = 0.001). *Peak* also differed significantly from *Non-Peak* in CGRP-injected mice for both Facial Grimace Score (paired t-test, *Peak* vs. *Non-Peak* median difference = −21.58 percentage points, p < 0.05) and Body Position Score (paired t-test, *Peak* vs. *Non-Peak* median difference = −47.1 percentage points, p < 0.05). These data indicate that mouse pain behavior during the empirical *Peak* is distinct from that during *Non-Peak* periods. The *Non-Peak* and *Peak* median values were therefore used to create thresholds for two levels of pain, as indicated in Figures 5A-B: Level 1 (dotted line; −19.04 percentage points difference from *Baseline* for facial grimace and −132.4 percentage points for body position) and Level 2 (dashed line; −40.62 percentage points difference from *Baseline* for facial grimace and −179.5 percentage points for body position).

Selection of the empirical peak which led to the use of *Peak* and *Non-Peak* periods throughout this study (starting with Figure 2) was based on diary plots binned into one-minute segments across the entire CGRP recording period (Figure 5C-D). Scores for each one-minute segment were the summed value of all individual measurements as a median percent difference from *Baseline*. It was found that from minute 27 to 32 the binned scores were below the mean value of *All* CGRP frames in Facial Grimace Score (−27.17 percentage points, Figure 5C, grey bars), and from minute 26 to 33 they were below the mean value of *All* CGRP frames in Body Position Scores (−145.7 percentage points, Figure 5D, grey bars). The empirical *Peak* was therefore selected based on the criteria where both scores were below the mean of *All* CGRP frames for both scores. It is noteworthy that other one-minute binned values outside of the empirical peak fall below the mean from the *All* CGRP recording (Figure 5C-D, grey bars), however, the longest continuous period where both scores meet these criteria was used as the empirical *Peak*. This method confirmed the previously reported occurrence of the peak of CGRP-induced migraine pain approximately 30 minutes after injection (24). Due to a higher sampling rate and continuous recording in the present study (30 frames/s, 20 minutes recording time), the empirical *Peak* was found to span 5 minutes and was used to define a second pain level.

In summary, automated analysis of video recordings of mouse behavior using both Facial Grimace and Body Position Scores provides a high throughput approach to quantify persistent pain in mice. This method was built on established methods for detecting facial grimace (23) and combined this with body position changes using a single camera in the same chamber. Validation of this method using CGRP-induced migraine demonstrated the ability to detect persistent pain and further allowed for the defining of two migraine pain thresholds (Levels 1 and 2).

### Behavioral detection of sound-evoked pain

Having validated the above method to measure persistent pain from video recordings of mouse behavior, this approach was next used to detect changes in behavior elicited by prolonged sound exposure. Sensitivity to sound in mice has been detected primarily using behavior measurements in response to sound onset (4, 15, 16), which is an effective way to monitor changes in sound sensitivity. To measure behavioral changes associated with pain caused by longer duration sound, the following experiments used facial grimace and body position measurements to quantify sound-evoked pain and determine the level of sound that caused pain-related behaviors in mice.

### Sound exposure elicits changes in individual measurements of facial grimace and body position

Mouse behavior was next tested during a sound exposure protocol with varying sound intensities using an assessment of facial grimace and body position parameters as described above. Individual facial grimace and body position parameters changed with increasing sound intensities (Figure 6). The experimental design is shown in Figure 6A. Video recordings started 8 minutes after the mice were placed in the acoustic chamber. The recording from 8-10 minutes was used as the baseline recording. Sound exposures of varied sound intensity levels in a set sequence were performed in 2-minute time-periods and were interleaved with 2-minute time-periods of no sound exposure (‘*No Sound*’). The highest sound exposure level (120 dB) was placed last in the sequence of exposures, assuming that lasting cochlear damage might occur.

**Figure 6.**
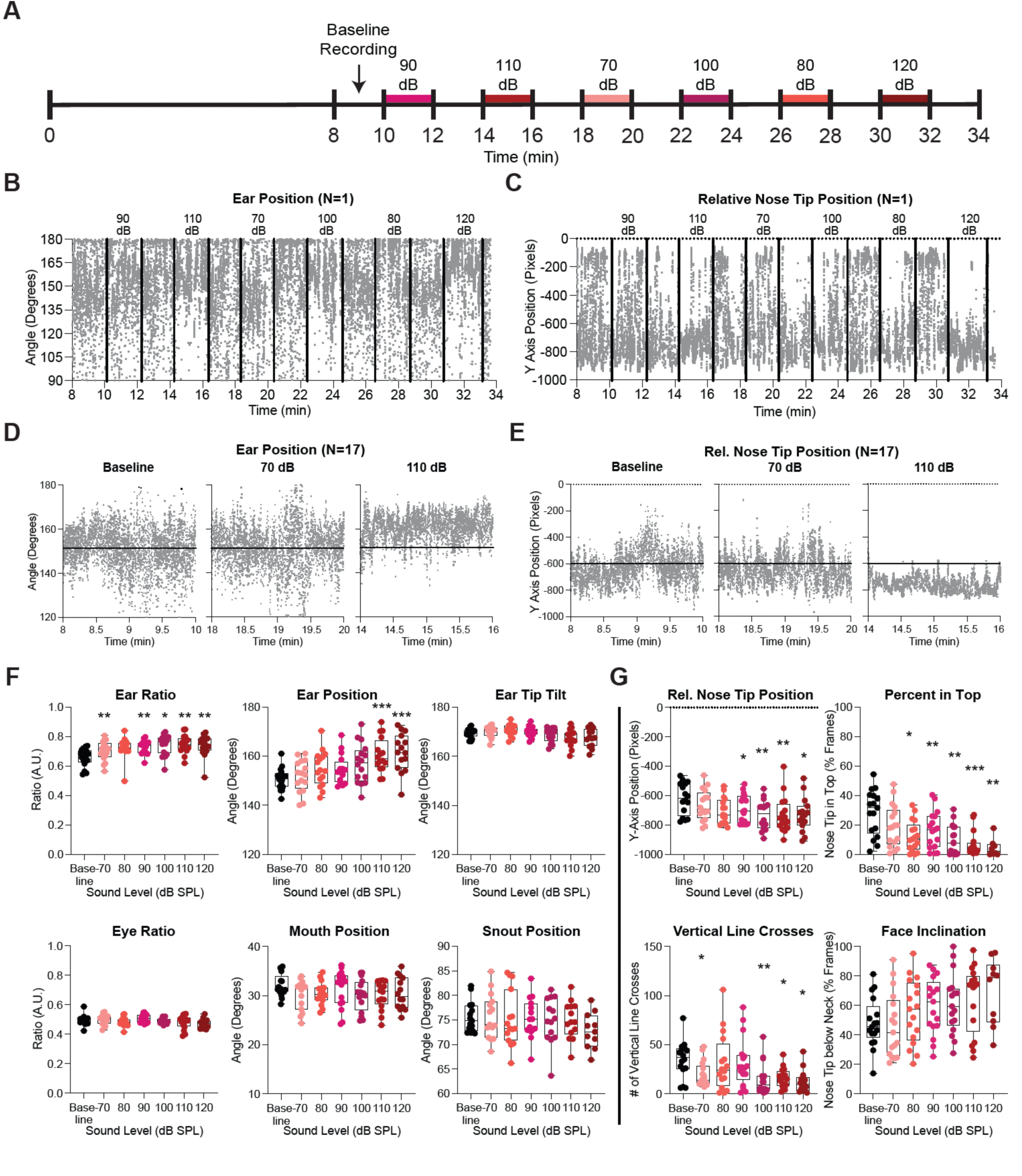
Sound exposure elicits changes in individual measurements of facial grimace and body position. **A**) Timeline of interleaved sound level experiments. Recording started at 8 minutes after the mice were placed in the acoustic chamber. The recording from 8-10 minutes was used as the baseline recording. Sound exposure was initiated in 2-minute time-periods separated by 2 minutes of silence. Sound was presented in out of decibel-level order as indicated by color and label. **B**) Example of analyzed frames of Ear Position measurements for one mouse for the entirety of one recording (N=1). The black vertical lines indicate when the sound levels were manually changed on or off. **C**) Example of analyzed frames of Relative Nose Tip Position measurements for one mouse for the entirety of one recording (N=1). The black vertical lines indicate when the sound levels were manually changed on or off. **D**) Averaged Ear Position measurements of analyzed frames from 12 mice (N=12) for Baseline, 70 dB, and 110 dB time periods. The time in minutes is the average on and off time periods for Baseline, 70 dB and 110 dB for each recording. The 70 dB recording occurred after the 110 dB recording, but the graphs have been reordered according to decibel level. Horizontal black lines indicate the average for the baseline graph. **E**) Averaged Relative Nose Tip Position measurements of analyzed frames from 12 mice (N=12) for Baseline, 70 dB, and 110 dB time periods. The time in minutes is the average on and off time periods for Baseline, 70 dB and 110 dB for each recording. The 70 dB recording occurred after the 110 dB recording, but the graphs have been reordered according to decibel level. Horizontal black lines indicate the average for the baseline graph. **F**) The following graphs show the averaged data for each facial parameter indicated. Each dot is the averaged data for the indicated 2-minute time-period for 1 mouse. The sound levels have been reordered from lowest to most intense sound level. Box and whiskers have a line at the median and the box is the interquartile range with the whiskers extending to the maximum and minimum value of the data. Ear Ratio: baseline compared to each sound level, repeated measures mixed model, p=0.0013; post hoc comparison using Tukey’s correction, baseline compared to 70 dB,**, p=0.0068, baseline compared to 90 dB, **, p=0.0055, baseline compared to 100 dB, *, p=0.0135, baseline compared to 110 dB, **, p=0.0079, baseline compared to 120 dB, **, p=0.0015. Ear Position: baseline compared to each sound level, repeated measures mixed model, p=<0.0001; post hoc comparison using Tukey’s correction, baseline compared to 110 dB, ***, p=0.0001, baseline compared to 120 dB, ***, p=0.0008. Ear Tip Tilt: baseline compared to each sound level, repeated measures mixed model, p=0.0148; Eye Ratio: baseline compared to each sound level, repeated measures mixed model, p=0.0354; Mouth Position: baseline compared to each sound level, repeated measures mixed model, ns, p=0.1308. Snout Position: baseline compared to each sound level, repeated measures mixed model, ns, p=0.5743. **G**) The following graphs show the averaged data for each body position parameter indicated by the title. Each dot is the averaged data for the indicated 2-minute time-period for 1 mouse. The sound levels have been reordered from lowest to most intense sound level. Box and whiskers have a line at the median and the box is the interquartile range with the whiskers extending to the maximum and minimum value of the data. Rel. Nose Tip Position: baseline compared to each sound level, repeated measures mixed model, p=0.0016; post hoc comparison using Tukey’s correction, baseline compared to 90 dB, *, p=0.0142, baseline compared to 100 dB, **, p=0.0095, baseline compared to 110 dB, **, p=0.0058, baseline compared to 120 dB, *, p=0.0473. Percent in Top: baseline compared to each sound level, repeated measures mixed model, p=<0.0001; post hoc comparison using Tukey’s correction, baseline compared to 80 dB, *, p=0.0186, baseline compared to 90 dB, **, p=0.0022, baseline compared to 100 dB, **, p=0.0026, baseline compared to 110 dB, ***, p=0.0005, baseline compared to 120 dB, **, p=0.0066. Vertical Line Crosses: baseline compared to each sound level, repeated measures mixed model, p=0.0078; post hoc comparison using Tukey’s correction, baseline compared to 70 dB, *, p=0.0112, baseline compared to 100 dB, **, p=0.0018, baseline compared to 110 dB, *, p=0.0122, baseline compared to 120 dB, *, p=0.0122. Face Inclination: baseline compared to each sound level, repeated measures mixed model, p=0.0221.

Example diary plots of one mouse show individual measures of Ear Position and Relative Nose Tip Position in response to the sound exposure protocol (Figure 6B-C). Superimposed data from all mice (n = 17) visualize the results from *Baseline*, 70 dB SPL, and 110 dB SPL periods (Figure 6D-E, raw and labeled video examples shown in Supplemental Video 2 and 3). Median values along with appropriate statistical tests revealed differences compared with *Baseline* in multiple individual measurements of facial grimace and body position at many intensities of sound. Ear Ratio, Ear Position, Relative Nose Tip Position, Percent in Top, and Vertical Line Crosses were all significantly different at 110 dB SPL compared with Baseline (Figure 6F-G). Specifically, Ear Ratio (repeated measures mixed effects model of all groups, p = 0.0013, post-hoc comparisons using Tukey’s correction, 70 dB vs. *Baseline* median difference = 0.0361 AU, p < 0.01, 90 dB vs. *Baseline* median difference = 0.0602 AU, p < 0.01, 100 dB vs. *Baseline* median difference = 0.0927 AU, p < 0.05, 110 dB vs. *Baseline* median difference = 0.0637 AU, p < 0.01, and 120 dB vs. *Baseline* median difference = 0.0642 AU, p < 0.01), Ear Position (repeated measures mixed effects model of all groups, p < 0.0001, post hoc comparisons using Tukey’s correction, 110 dB vs. *Baseline* median difference = 8.0°, p < 0.001, 120 dB vs. *Baseline* mean difference = 10.6°, p < 0.001), Relative Nose Tip Position (repeated measures mixed effects model of all groups, p = 0.0016, post hoc comparisons using Tukey’s correction, 90 dB vs. *Baseline* median difference = − 87 pixels, p < 0.05, 100 dB vs. *Baseline* median difference = −106 pixels, p < 0.01, 110 dB vs. *Baseline* median difference = −148 pixel, p < 0.01, 120 dB vs. *Baseline* median difference = −114 pixels, p < 0.05), Percent in Top (repeated measures mixed effects model of all groups, p < 0.0001, post hoc comparisons using Tukey’s correction, 80 dB vs. *Baseline* median difference = −21.4 percentage points, p < 0.05, 90 dB vs. *Baseline* median difference = −15.0 percentage points, p < 0.01, 100 dB vs. *Baseline* median difference = −24.1 percentage points, p < 0.01, 110 dB vs. *Baseline* median difference = −28.9 percentage points, p < 0.001, 120 dB vs. *Baseline* median difference = −29.4 percentage points, p < 0.01), and Vertical Line Crosses (repeated measures mixed effects model of all groups, p = 0.0078, post hoc comparisons using Tukey’s correction, 70 dB vs. *Baseline* median difference = −23 incidences, p < 0.05, 100 dB vs. *Baseline* median difference = −28 incidences, p < 0.01, 110 dB vs. *Baseline* median difference = −22 incidences, p < 0.05, 120 dB vs. *Baseline* median difference = −27 incidences, p < 0.05) all showed significant differences at multiple intensities. Ear Tip Tilt, Eye Ratio, and Face Inclination were all significant in the mixed effects model but showed no significant post-hoc comparisons. Specifically, Ear Tip Tilt (repeated measures mixed effects model of all groups, p **=** 0.0148), Eye Ratio (repeated measures mixed effects model of all groups, p = 0.0354), and Face Inclination (repeated measures mixed effects model of all groups, p = 0.0221) showed significant group effects. Finally, Mouth and Snout Position measures had no significant group effects. Taken together, these data suggest that sound-evoked pain may be preferentially detected by individual grimace measurements related to ears, and by changes in body position.

### After individual sound exposures, mouse behavior returned to baseline during *No Sound* periods except after exposure to sound at 120 dB SPL

The sound exposure protocol for individual mice included a sequence of multiple sound presentations with varying sound pressure levels, and the analysis of the resulting data, as presented in Figure 6, relies on the assumption that mouse behavior returns to baseline following each sound presentation. To verify this assumption, *No Sound* periods following each sound presentation were analyzed. In Figure 7, *No Sound* periods are arranged in increasing order of the preceding sound exposure level, and not in the order as they were presented in the interleaved sound exposure protocol. Ear ratio showed a significant effect of group (Figure 7A; repeated measures mixed effects model, p = 0.0012), however no significant differences were detected in post hoc comparisons to *Baseline* in any *No Sound* period. The remaining facial grimace parameters showed no significant changes following any sound intensity presented compared to *Baseline*. By contrast, for most body position measurements, the *No Sound* period after 120 dB SPL sound exposure did show significant differences compared with *Baseline* (Figure 7B).

**Figure 7.**
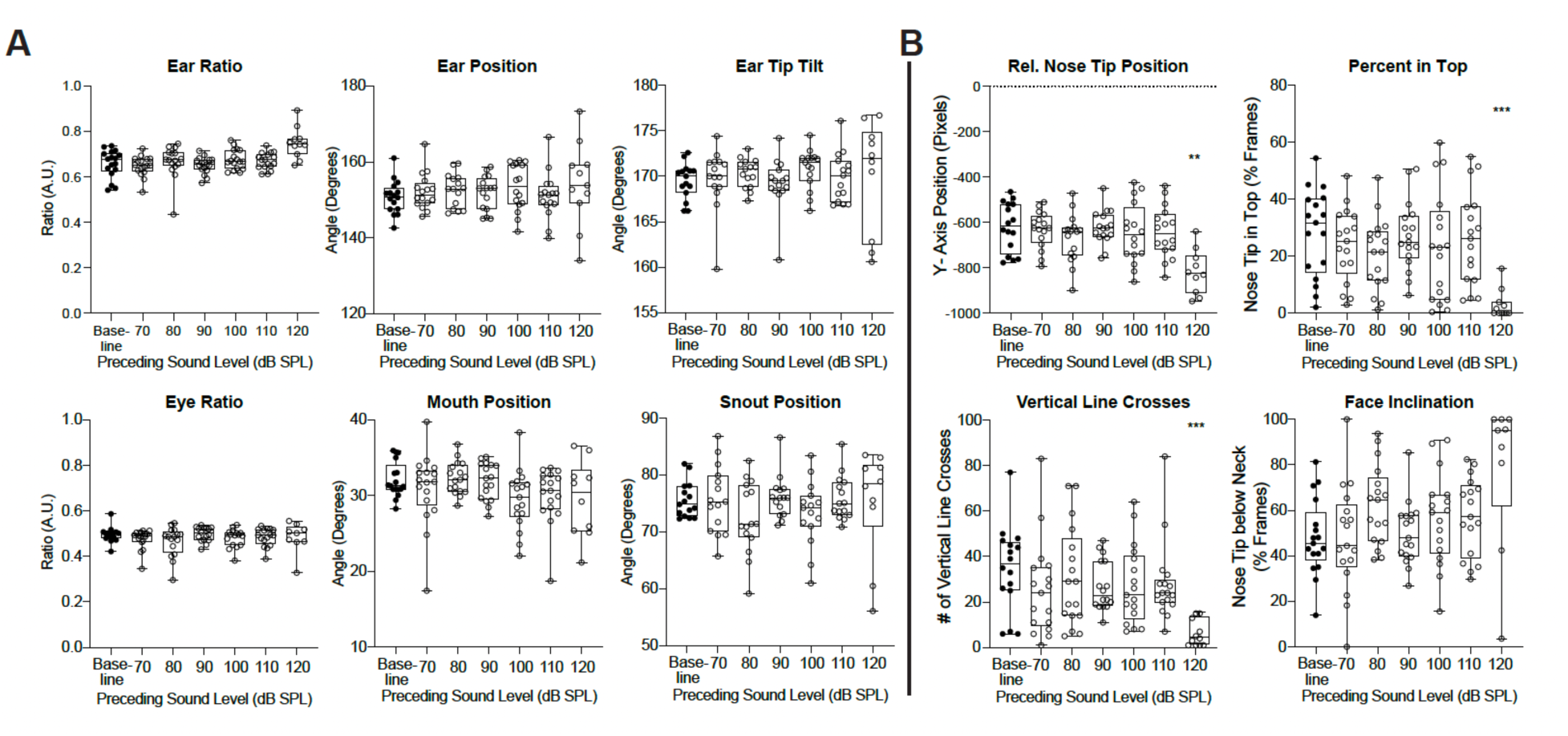
After sound exposures, mouse behavior returned to baseline during *No Soun* periods except after exposure to sound at 120 dB SPL. A) The graphs show the averaged data for each facial grimace parameter indicated by the title. Each dot is the averaged data for the indicated 2-minute time-period for 1 mouse. The data have been reordered based on the preceding sound level of the no sound period from lowest to most intense sound level. Box and whiskers have a line at the median and the box is the interquartile range with the whiskers extending to the maximum and minimum value of the data. Ear Ratio: baseline compared to each sound level, repeated measures mixed model, p=0.0012. All other facial parameters showed no significant changes. **B**) The graphs show the averaged data for each body position parameter indicated by the title. Each dot is the averaged data for the indicated 2-minute time-period for 1 mouse. The data have been reordered based on the preceding sound level of the silent sound period from lowest to most intense sound level. Box and whiskers have a line at the median and the box is the interquartile range with the whiskers extending to the maximum and minimum value of the data. Rel. Nose Tip Position: baseline compared to each sound level, repeated measures mixed model, p=0.0003; post hoc comparison using Tukey’s correction, baseline compared to 120 dB, **, p=0.002. Percent in Top: baseline compared to each sound level, repeated measures mixed model, p=0.0013; post hoc comparison using Tukey’s correction, baseline compared to 120 dB, ***, p=0.001. Vertical Line Crosses: baseline compared to each sound level, repeated measures mixed model, p=0.005; post hoc comparison using Tukey’s correction, baseline compared to 120 dB, ***, p=0.0001. Face Inclination: baseline compared to each sound level, repeated measures mixed model, p=0.0161.

Specifically, Relative Nose Tip Position (repeated measures mixed effects model, p = 0.0003, post-hoc comparisons using Tukey’s correction, post-120 dB vs. *Baseline* median difference = −207 pixels, p < 0.01), Percent in Top (repeated measures mixed effects model, p = 0.0013, post-hoc comparisons using Tukey’s correction, post-120 dB vs. *Baseline* median difference = −32 percentage points, p < 0.001), and Vertical Line Crosses (repeated measures mixed effects model, p = 0.01, post-hoc comparisons using Tukey’s correction, post-120 dB vs *Baseline* median difference = −33 incidences, p < 0.001) all showed significant changes during *No Sound* following 120 dB SPL, while no significant differences were found in post-hoc comparisons for Face Inclination (repeated measures mixed effects model, p = 0.0161). In this current study, 120 dB SPL was presented last with the consideration that this level of sound might impact subsequent animal behavior, which was confirmed here for multiple individual body position measurements.

In Summary, these findings indicate that for the specific design of the sound protocol used here, after sound exposures mouse behavior returned to baseline during *No Sound* periods, except after exposure to sound at 120 dB SPL.

### Sound exposure above 100 dB SPL caused changes in behavior consistent with pain in mice

As presented for the migraine model (Figure 5), Facial Grimace and Body Position Scores for both *Sound* and *No Sound* were calculated as a sum of the changes from *Baseline* in individual measurements (Supplemental Figure 3) (Figure 8). For comparison, thresholds for two migraine pain levels, Level 1 and Level 2, as established in the migraine model (Figure 5), were added to Figure 8. Both composite scores for ‘Sound’ were significantly different from *Baseline* for sound presentation at 100 dB SPL and above. Additionally, median values of both composite scores at 100 dB SPL and above were found to surpass the migraine pain Level 1 threshold (Figure 8A; dotted line). This validates that both stimuli, sound and migraine, elicit significant behavioral changes from *Baseline*, surpassing a threshold for pain to a similar extent. Specifically, significant differences were found for Grimace Scores (repeated measures mixed effects model, p < 0.0001, post-hoc comparisons using Tukey’s correction, 100 dB vs *Baseline* median difference = −22.6 percentage points, p < 0.01, 110 dB vs. *Baseline* median difference = −30.9 percentage points, p < 0.0001, and 120 dB vs. Baseline median difference = −29.6 percentage points, p < 0.001) and Position Scores (repeated measures mixed effects model, p = 0.0012, post-hoc comparisons using Tukey’s correction, 100 dB vs. *Baseline* median difference = −165.1 percentage points, p < 0.001, 110 dB vs *Baseline* median difference = −141.9 percentage points, p < 0.05, 120 dB vs Baseline median difference = −98.6 percentage points, p < 0.001) at and above 100 dB SPL. These findings indicated that mice exposed to 2 – 20 kHz sound for two minutes at intensities of 100 dB SPL and greater showed behavioral signatures consistent with pain.

**Figure 8.**
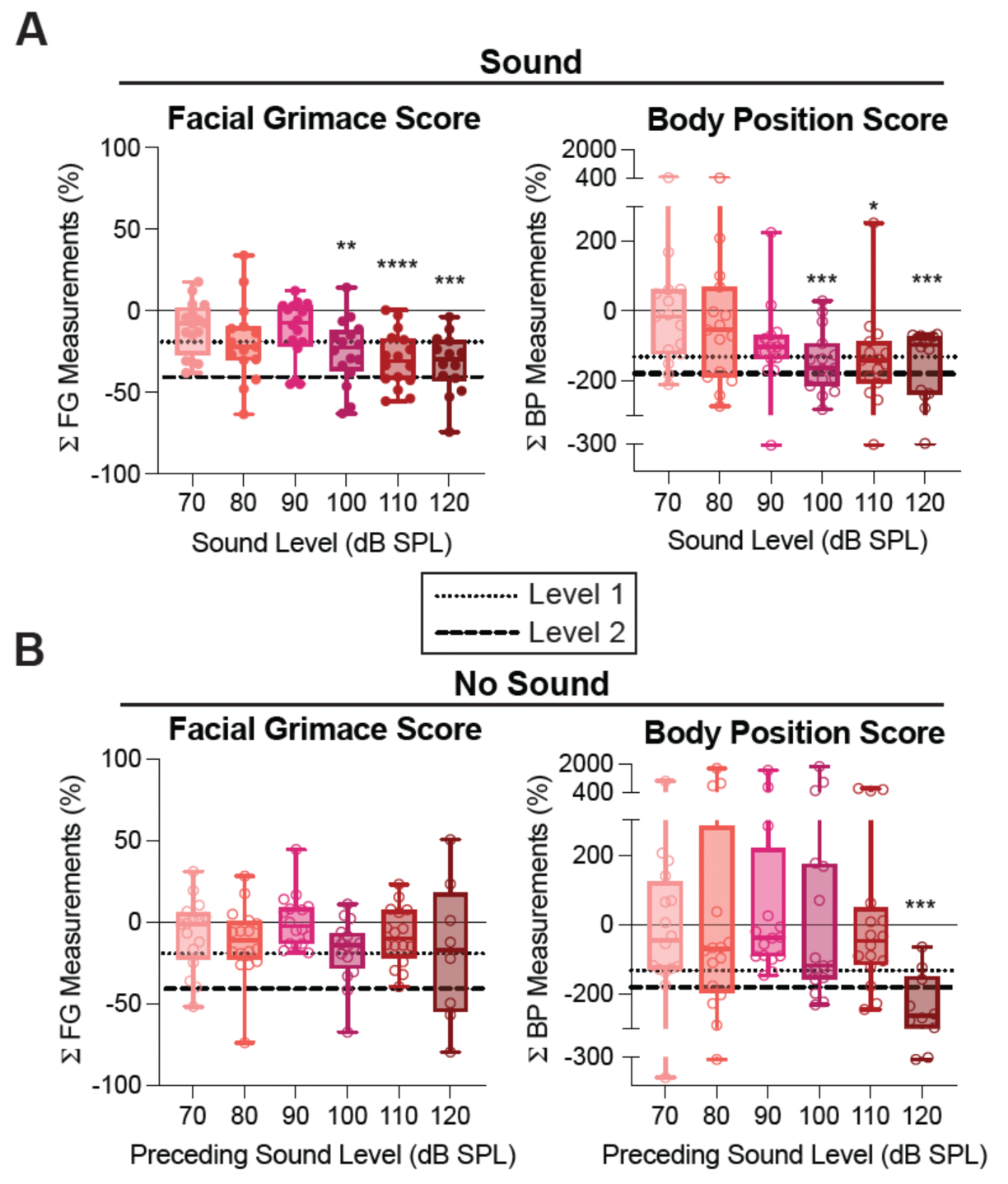
Sound exposure above 100 dB SPL caused changes in behavior consistent with pain in mice. **A**) Facial Grimace (left) and Body Position (Right) Scores for mice during exposure to sound (N=17). Facial Grimace Score, Sound: repeated measures mixed effects model, p=<0.0001; post hoc comparisons using Tukey’s correction; baseline compared to 100 dB, **, p=0.0025, baseline compared to 110 dB, ****, p=<0.0001, baseline compared to 120 dB, ***, p=0.0003. Body Position Score, Sound: repeated measures mixed effects model, p=0.0012; post hoc comparisons using Tukey’s correction; baseline compared to 100 dB, ***, p=0.0003, baseline compared to 110 dB, *, p=0.0174, baseline compared to 120 dB, ***, p=0.0006. The Pain Level 1 threshold is − 19.04 for Facial Grimace and −132.4 for Body Position (dotted lines). The Pain Level 2 threshold is −40.62 for Facial Grimace and −179.5 for Body Position (dashed lines). **B**) Facial Grimace (Left) and Body Position (Right) Scores for mice during the no sound periods (N=17). Box and whiskers have a line at the median and the box is the interquartile range with the whiskers extending to the maximum and minimum value of the data. Facial Grimace Score, No Sound: baseline compared to each sound level, repeated measures mixed model, ns, p=0.1157. Body Position Score, No Sound: baseline compared to each sound level, repeated measures mixed model, ns, p=0.2341

Composite scores for *No Sound* had no significant effects from the mixed model analyses for facial grimace or body position (Figure 8B). It is noteworthy that during the period of no sound following 120 dB SPL, the Body Position Score surpassed the migraine pain Level 2 threshold (Figure 8B; dashed line). This change in behavior following 120 dB SPL is consistent with statistically significant changes observed in individual measurements described above (Figure 7 and Supplemental Figure 3).

### The contribution of cochlear mechano-transduction to generating sound-evoked pain behavior

Pain caused by loud sounds could result from stress forces on auditory structures peripheral to the cochlea, such as sound pressure activating stretch receptors in the tympanic membrane or middle ear (44). To test whether normal cochlear function is necessary for sound-evoked pain, mice that lack mechano-transduction due to the loss the TMIE protein needed for proper function of the mechano-transduction channel in all cochlear hair cells (*Tmie*-knockout mice, *Tmie^-/-^*) and littermate heterozygous or wild-type controls (*Tmie^CTL^*) were presented with the same interleaved sound protocol described above and tested for pain behavior (27–30). Importantly, both heterozygous and wild-type mice show normal cochlear mechano-transduction and thus can serve as valid controls (45). These mice are maintained on a C57BL/6J background.

### Mice lacking TMIE largely showed no changes in individual behavioral measurements in response to sound exposure

Individual measurements of facial grimace and body position in *Tmie^CTL^*mice (Figure 9) showed similar trends to those of the wild-type C57BL/6J mice described above. Example diary plots from an individual *Tmie^CTL^* mouse demonstrate changes in Ear Position and Relative Nose Tip Position during sound exposures (Figure 9A-B). For data from all mice combined, for facial grimace measurements, Ear Ratio and Ear Position were significantly different from baseline at multiple sound levels (Figure 9C). Specifically, significant increases were found in Ear Ratio (repeated measures mixed effects model, p = 0.0009, post-hoc comparisons using Tukey’s correction, 90 dB vs. Baseline median difference = 0.127 AU, p < 0.001, 100 dB vs. Baseline median difference = 0.173 AU, p < 0.01, and 120 dB vs. Baseline median difference = 0.117 AU, p < 0.05) and in Ear Position (repeated measures mixed effects model, p = 0.0245, post-hoc comparisons using Tukey’s correction, 110 dB vs. Baseline median difference = 3.1°, p < 0.05, and 120 dB vs. Baseline median difference = 16.5°, p < 0.01). Eye ratio showed a significant group effect but no post-hoc tests were significantly different from baseline (repeated measures mixed effects model, p = 0.0293). For body position measurements, significant reductions were found in Relative Nose Tip Position, Percent in Top, Vertical Line Crosses (Figure 9D). Specifically, Relative Nose Tip Position (repeated measures mixed effects model, p = 0.014, post-hoc comparisons using Tukey’s correction, 110 dB vs. Baseline median difference = −129 pixels, p < 0.01), Percent in Top (repeated measures mixed effects model, p = 0.0124, post-hoc comparisons using Tukey’s correction, 110 dB vs. Baseline median difference = −15.1%, p < 0.05), Vertical Line Crosses (repeated measures mixed effects model, p = 0.0078, post-hoc comparisons using Tukey’s correction, 90 dB vs. Baseline median difference = −51 incidences, p < 0.05, 100 dB vs. Baseline median difference = −53 incidences, p < 0.01, 110 dB vs. Baseline median difference = −60 incidences, p < 0.01) all showed significant decreases at 110 dB SPL (e.g., see Supplemental Video 4).

**Figure 9.**
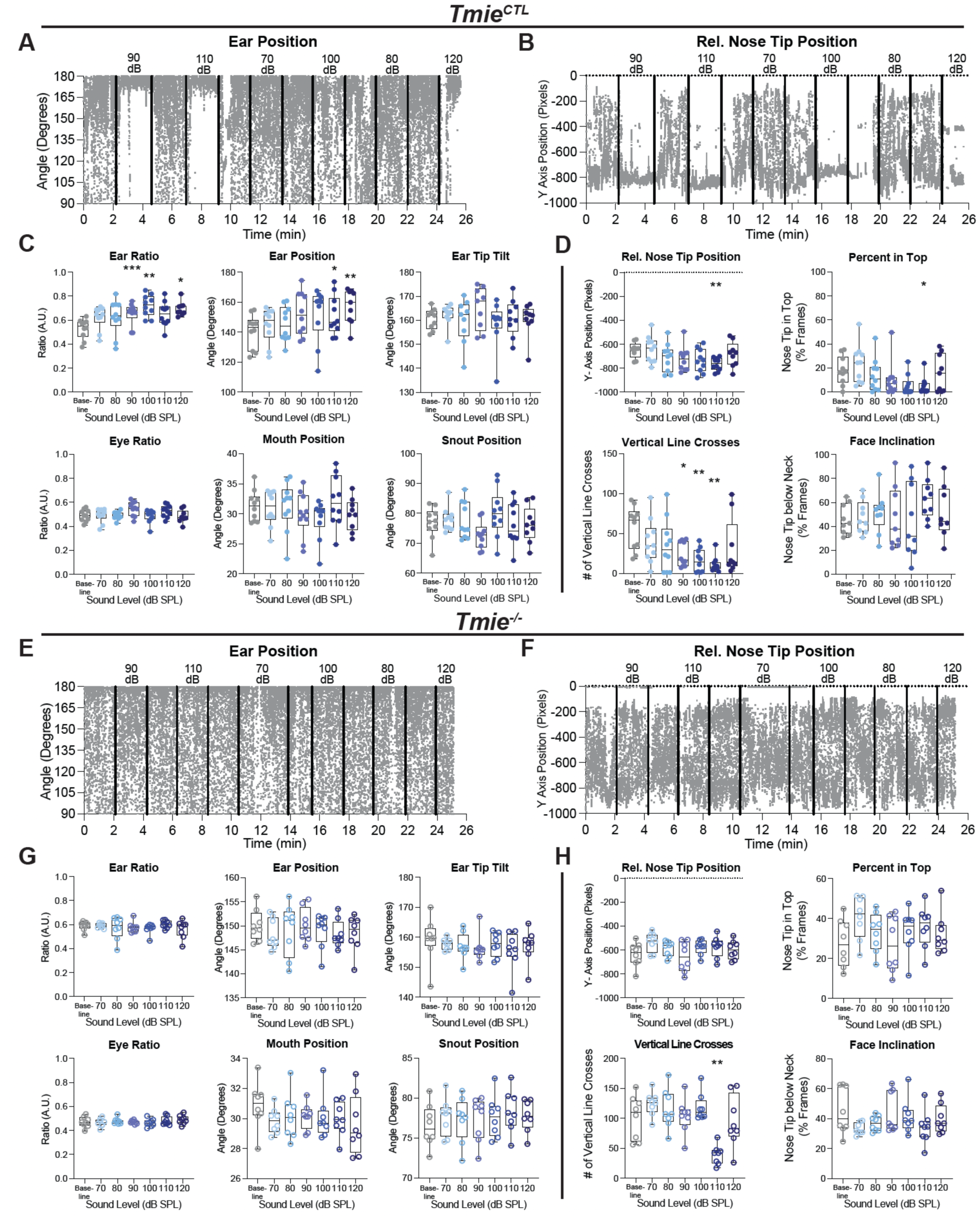
Mice lacking TMIE largely showed no changes in individual behavioral measurements in response to sound exposure. **A**) Example of analyzed frames of Ear Position measurements for one *Tmie^CTL^* mouse for the entirety of one recording. The black vertical lines indicate when the sound levels were manually changed on or off. **B**) Example of analyzed frames of Rel. Nose Tip Position measurements for one *Tmie^CTL^* mouse for the entirety of one recording. The black vertical lines indicate when the sound levels were manually changed on or off. **C**) The following graphs show the averaged data for each facial parameter indicated. Each dot is the averaged data for the indicated 2-minute time-period for 1 *Tmie^CTL^* mouse (N=10). The sound levels have been reordered from lowest to most intense sound level. Box and whiskers have a line at the median and the box is the interquartile range with the whiskers extending to the maximum and minimum value of the data. Ear Ratio: repeated measures mixed effects model, p=0.0009; post hoc comparisons using Tukey’s correction, baseline compared to 90 dB, ***, p=0.0009, baseline compared to 100 dB, **, p=0.0049, baseline compared to 120 dB, *, p=0.0272. Ear Position: repeated measures mixed effects model, p=0.0245; post hoc comparisons using Tukey’s correction, baseline compared to 110 dB, *, p=0.027, baseline compared to 120 dB, **, p=0.0033. Eye Ratio: repeated measures mixed effects model, p=0.0293. **D**) The following graphs show the averaged data for each body position parameter indicated by the title. Each dot is the averaged data for the indicated 2-minute time-period for 1 *Tmie^CTL^* mouse (N=10). The sound levels have been reordered from lowest to most intense sound level. Box and whiskers have a line at the median and the box is the interquartile range with the whiskers extending to the maximum and minimum value of the data. Rel. Nose Tip Position: repeated measures mixed effects model, p=0.014; post hoc comparisons using Tukey’s correction, baseline compared to 110 dB, **, p=0.037. Percent in Top: repeated measures mixed effects model, p=0.0124; post hoc comparisons using Tukey’s correction, baseline compared to 110 dB, *, p=0.0224. Vertical Line Crosses: repeated measures mixed effects model, p=0.0078; post hoc comparisons using Tukey’s correction, baseline compared to 90 dB, *, p=0.0207, baseline compared to 100 dB, **, p=0.0026, baseline compared to 110 dB, **, p=0.0027. Face Inclination: repeated measures mixed effects model, ns, p=0.1787; post hoc comparisons using Tukey’s correction, baseline compared to 110 dB, *, p=0.0316. **E**) Example of analyzed frames of Ear Position measurements for one *Tmie^-/-^* mouse for the entirety of one recording. The black vertical lines indicate when the sound levels were manually changed on or off. **F**) Example of analyzed frames of Rel. Nose Tip Position measurements for one *Tmie^-/-^* mouse for the entirety of one recording. The black vertical lines indicate when the sound levels were manually changed on or off. **G**) The following graphs show the averaged data for each facial parameter indicated. Each dot is the averaged data for the indicated 2-minute time-period for 1 *Tmie^-/-^*mouse (N=8). The sound levels have been reordered from lowest to most intense sound level. Box and whiskers have a line at the median and the box is the interquartile range with the whiskers extending to the maximum and minimum value of the data. **H**) The following graphs show the averaged data for each body position parameter indicated by the title. Each dot is the averaged data for the indicated 2-minute time-period for 1 *Tmie^-/-^* mouse (N=8). The sound levels have been reordered from lowest to most intense sound level. Box and whiskers have a line at the median and the box is the interquartile range with the whiskers extending to the maximum and minimum value of the data. Rel. Nose Tip Position: repeated measures mixed effects model, p=0.0401. Note this relationship shows an increase Rel. Nose Tip Position not associated with pain. Vertical Line Crosses: repeated measures mixed effects model, p=0.0005; post hoc comparisons using Tukey’s correction, baseline compared to 110 dB, **, p=0.0028.

In contrast, in *Tmie^-/-^* mice, which lack functional cochlear mechano-transduction, most individual facial grimace measurements were not significantly different from baseline in any condition, except for one body position measurement that did show significant differences vs. baseline (detailed below). Example diary plots of Ear Position and Relative Nose Tip Position from an individual mouse show no obvious behavioral changes during any sound exposure (Figure 9E-F). For data from all mice combined, individual facial grimace measurements did not show any changes (Figure 9G). However, of the four body position measurements, only Vertical Line Crosses were significantly reduced vs. baseline at 110 dB SPL (Figure 9H; repeated measures mixed effects model, p = 0.0005, post-hoc comparisons using Tukey’s correction, 110 dB vs. Baseline mean difference = −68 incidences, p < 0.01). It is unclear what could cause the decreased turning observed here, however it is noteworthy that *Tmie*^-/-^ mice have altered vestibular function that causes repeated turning behavior, which caused a two-fold increase in Vertical Line Crosses at baseline vs. *Tmie^CTL^* mice (see Figure 9D and H). Relative Nose Tip Position also showed a significant group effect (repeated measures mixed effects model, p = 0.04), but the direction of this effect was towards increased position and thus was not suggestive of pain.

### Cochlear mechano-transduction is required to generate sound-evoked pain behavior

Individual measurements of both facial grimace and body position were converted into percentage change in baseline (Supplemental Figure 4) and summed to generate composite scores (Figure 10). Facial Grimace and Body Position Scores revealed sound-evoked pain in *Tmie^CTL^* mice emerging at 100 dB SPL and greater sound intensities (Figure 10A) replicating the findings from C57BL/6J detailed above. Specifically, Facial Grimace Score and Body Position Score both showed significant changes at 100 and 110 dB SPL consistent with normal hearing C57BL/6J mice presented above (Figure 5) (Facial Grimace Score: repeated measures mixed effects model, p = 0.0004, post-hoc comparisons using Tukey’s correction, 90 dB vs. Baseline median difference = − 26.0%, p < 0.05, 100 dB vs. Baseline median difference = −53.7%, p < 0.01, 120 dB vs. Baseline median difference = −47.9%, p < 0.001; Body Position Score: repeated measures mixed effects model, p = 0.0031, post-hoc comparisons using Tukey’s correction, 100 dB vs. Baseline median difference = −154.8%, p < 0.001, 110 dB vs. Baseline median difference = −140%, p < 0.0001).

**Figure 10.**
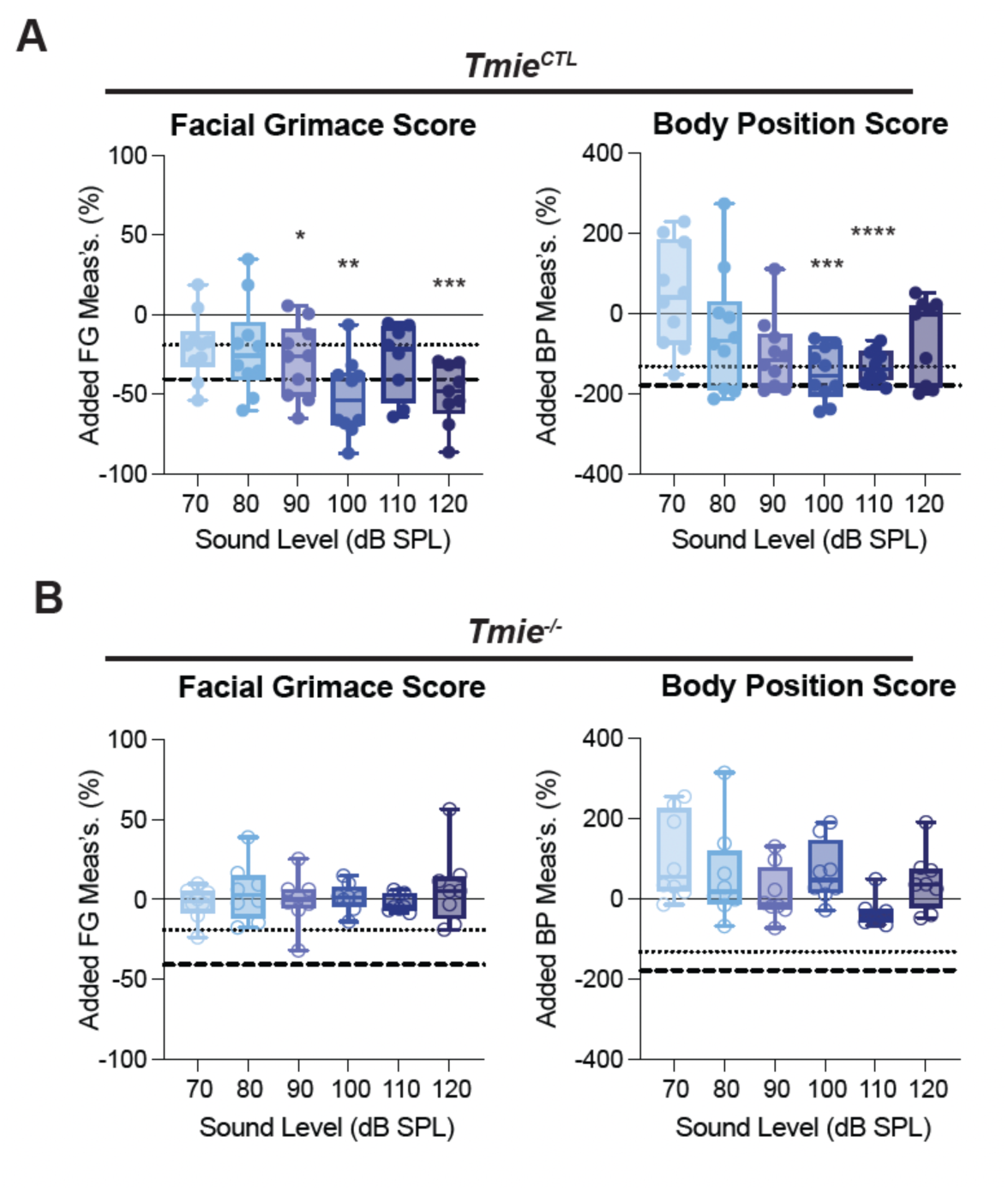
Cochlear mechano-transduction is required to generate sound-evoked pain behavior. Mice deficient in mechanotransduction do not show pain-associated behavioral changes when exposed to loud sounds like wildtype mice. **A**) Facial Grimace (Left) and Body Position (Right) Scores for *Tmie^CTL^* mice exposed to sound(N=10). Box and whiskers have a line at the median and the box is the interquartile range with the whiskers extending to the maximum and minimum value of the data. Facial Grimace Score, *Tmie^CTL^*: repeated measures mixed effects model, p=0.0004; post hoc comparisons using Tukey’s correction, baseline compared to 90 dB, *, p=0.0417, baseline compared to 100 dB, **, p=0.0011, baseline compared to 120 dB, ***, p=0.0007. Body Position Score, *Tmie^CTL^*: repeated measures mixed effects model, p=0.0031; post hoc comparisons using Tukey’s correction, baseline compared to 100 dB, ***, p=0.0007, baseline compared to 110 dB, ****, p=<0.0001. **B**) Facial Grimace (left) and Body Position (Right) Scores for *Tmie^-/-^* mice exposed to sound (N=8). Facial Grimace Score, *Tmie^-/-^* : repeated measures mixed effects model, ns, p=0.535; Body Position Score, *Tmie^-/-^*: repeated measures mixed effects model, p=0.0296. For a mouse to be experiencing pain, the scores must be lower than −19.04 for Facial Grimace and −132.4 for Body Position (dotted line). For a mouse to experience Pain Level 2, the scores must be lower than −40.62 for Facial Grimace and −179.5 for Body Position (dashed line).

In *Tmie^-/-^* mice, no significant differences were observed for Facial Grimace or Body Positions Scores at any sound intensity (Figure 10B; Supplemental Video 5) suggesting that normal cochlear mechano-transduction is necessary for inducing sound-evoked pain. These results suggest that sound pressure causing the activation of stretch receptors in peripheral auditory structures was not the source of sound-evoked pain as measured and defined here (46–48). However, additional mechanisms that require normal cochlear function to generate subsequent pathophysiological responses cannot be ruled out, such as descending motor fibers innervating peripheral structures (such as cranial nerve V innervation of the tensor tympani), which could become hyperactive due to prior sound exposures and thereby could contribute to sound-evoked pain (46, 49, 50).

In summary, this study demonstrates the feasibility of using a single camera angle to measure facial grimace and additional positional information from video recordings of mice to detect behavioral changes associated with persistent pain. Validation of this approach with a known painful stimulus (CGRP-induced migraine) and its application to an experimental condition (loud sound), allowed for the investigation of sound-evoked pain in mice. Future studies could apply this method to other persistent pain conditions or to investigate further the sources and mechanisms responsible for pain caused by sound exposure.

## Discussion

To gain a better understanding how sound-evoked pain is generated and processed, it is crucial to implement methodology that allow for unbiased measuring of when sound becomes painful. Sound-evoked pain was quantified using a deep neural network model trained to automatically place points on the face and body of mice that were video recorded while being presented with different intensities of sound. Using this method, changes in facial grimace (21–23, 25, 26) and body position (51–55) were analyzed to create an inventory capable of detecting pain-related behaviors in mice. Calcitonin gene-related peptide (CGRP) injection to induce migraine in mice is a well-established persistent pain model (24) that was used to validate this method. CGRP-induced migraine caused detectable changes in behavior compared to baseline, supporting the approaches’ applicability to study other forms of persistent pain. Sounds presented at intensities of 100 dB and above resulted in significant pain behavioral changes compared to baseline that appeared in both facial grimace and body position composite scores but were not present in mice lacking cochlear mechano-transduction. Taken together, these findings demonstrate that machine learning-based detection of pain behaviors during persistent painful states, such as CGRP-induced migraine and sound-evoked pain, can be reliably measured in mice.

### Machine learning-based automated detection of pain-related behaviors

The approach presented above used video recordings of freely moving mice during painful stimuli, namely migraines and loud sound presentations, that were analyzed using a deep neural network model trained to automatically place points on the face and body of the mice. These points were used to calculate various individual parameters of facial grimace (Figure 1) and body position (Figure 3) that changed during painful stimuli (e.g., migraine) compared with baseline behavior and were plotted both as raw values (Figures 2 and 4) and as a normalized percentage change from baseline (Supplemental Figure 2A).

The body position measurements in this method served two primary purposes: (1) to provide an additional metric of movement that complements the facial grimace score, and (2) to demonstrate the possibilities of quantifying data from one to two x-y coordinate points using just one camera. By analyzing the y-axis of the nose tip, the average position could be calculated over a specified period (Relative Nose Tip Position). In this instance, when the animal moves up and down within the chamber, the average y-value is higher compared to when the animal remains stationary in a hunched posture (Fig. 4). Additionally, by introducing horizontal or vertical boundaries within the video frames, it was possible to quantify the average amount of time the animal spent in a specific region of the frame (e.g., Percent in Top) or the frequency of crossing a defined boundary (e.g., Vertical Line Crosses). Furthermore, the relationship between the y-values of two points could be analyzed to calculate the percentage of time this relationship was maintained (Face Inclination). These measurements provide examples of how pose estimation marker data can be utilized.

The percentage change from baseline for each facial grimace and body position parameter were separately summed to create two composite scores (Figure 5) to detect changes in behavior associated with CGRP-induced migraine pain. Since injections of CGRP intraperitoneally in mice produce a migraine with a well-documented time course (23), where the peak of the pain occurs approximately 30 minutes after injection, video recordings were performed from t = 20-40 minutes following injection. Strikingly, an empirical peak emerged from the analysis of these videos centered around the expected peak of 30 minutes, where different magnitudes of changes in pain-related behaviors were observed from t = 27-32 minutes compared to the remainder of the recording period before and after the peak. The median of each of these two periods, above referred to as “*Peak*” and “*Non-Peak*”, both differed significantly from baseline as well as from each other allowing for the delineation of two thresholds of migraine pain (Levels 1: *Non-Peak* and 2: *Peak*; Figure 5C-D). The ability of the approach to delineate two pain levels provides an advancement for pain measurement.

### Sound presentation evokes pain-related behaviors at and above 100 dB SPL

Humans can experience pain when exposed to loud sounds at 120 dB and above (56). However, the sound exposure levels and protocols that might cause loudness intolerances in laboratory mice are not well known. With that in mind, suprathreshold levels of sound from 70 dB to 120 dB were used to probe whether mice would display behavioral changes associated with pain at any of these levels of sound using the now validated method of machine learning-based detection of pain-related behaviors. Sound was presented for two-minute-long periods and periods of silence (‘*No Sound*’) occurred in between each sound presentation (Figure 6). The purpose of the ‘*No Sound*’ periods was to prevent an adaptation effect by continuous sound presentation and to serve as a control that behavioral changes restricted to sound presentation were reversible, which was true except for 120 dB SPL (see Figures 7 and 8). Both Facial Grimace and Body Position Scores were significantly changed from baseline starting at 100 dB SPL. It needs to be kept in mind, that the determined ‘pain thresholds’ are specific to the chosen experimental setting including the sound protocol and mouse model, and have to be reexamined whenever used under different conditions. The strength of the approach does not lie in the absolute measured threshold levels, but in the examination of a specific mouse model/condition, in comparison with the control conditions measured in the same settings.

### Probing auditory structures responsible for sound-evoked pain using *Tmie*^-/-^ mice

It remains unclear which peripheral or central auditory structures are necessary for sound-evoked pain. Multiple theories exist for the anatomical sources of sound-evoked pain (8, 13, 14), most of which posits that the process is initiated by pathological changes in the function of peripheral auditory structures that ultimately alter the function of central auditory processing, resulting in enhanced efferent activation of the tensor tympani muscle (44,50,58), hyperactivity of type II auditory nerve fibers which may act as auditory nociceptors (59–61), and/or hypoactivation of the auditory cortex and concurrent increased activity in the thalamus (10–12). However, it is conceivable that sound-evoked pain, especially when caused by loud sounds, may not rely on cochlear mechano-transduction in the inner ear, but rather could be caused by the activation of stretch receptors in the middle ear or on the tympanic membrane that could be triggered at high pressure levels to induce pain due to sound (46–48, 62).

*Tmie*^-/-^ mice (30–33, 45) exhibit normal development of outer and middle ear, but have profound inner ear sensory dysfunction, and were therefore here used to investigate the contributions of peripheral auditory structures to sound-evoked pain as measured using the method described above. *Tmie*^-/-^ mice showed no significant behavioral changes at any sound exposure level that reached the pain threshold for either the Facial Grimace or Body Position Scores (Figures 9 and 10). The littermate-control mice (*Tmie^CTL^*) exhibited similar behavioral changes associated with sound presentation as the wild-type C57BL/6J mice previously shown, indicating pain perception during loud sound exposure. Thus, cochlear function in *Tmie^CTL^* was necessary for a normally hearing animal to display sound-evoked pain behavior. It is still possible that in animals with disordered hearing or changes to the middle ear (such as otitis media), the middle ear could play a role in auditory pain perception. As in tinnitus and paralleled in phantom limb syndrome (63), central processing of auditory pain also likely plays a role in auditory pain disorders, such as pain hyperacusis (64), however, sound detection by the peripheral auditory system appears to be necessary for the generation of auditory pain in response to loud sounds in a normally hearing mouse.

### Implications for Animal Noise Exposure Protocols

Loud sound exposures are frequently used to study noise-induced hearing loss (NIHL) in animals. Specifically, loud sound protocols are used to induce temporary or permanent hearing damage to cochlear function as tested by ABR thresholds and morphological integrity of the cochlea (hair cell loss; synaptopathy) (65, 66). Such protocols often use 2 hours of 110 dB SPL (or greater) sound, exceeding the levels causing pain-behavior as reported here (61, 67). It is noteworthy that similar noise exposures have been shown to cause variable degrees of cochlear damage across institutions and mouse strains (68), suggesting that pain responses may also differ under these conditions. While intense noise levels are necessary to address specific research questions related to NIHL, the findings presented here highlight the possibility of additional behavioral effects accompanying such exposures.

### Expanding existing methods for detecting auditory-evoked behavior

Several methods exist to measure sound-evoked behavior in rodents. These methods fall into two categories: 1) measurements of reflexive responses to sound 2) measurements of aversion to sound. Acoustic startle responses and sub-second facial responses model loudness perception (12, 69), while conditioned aversion assays measure the preference of a mouse for a well-lit chamber over a dark chamber with noise going against the animal’s instinct (69, 70). Thus, loudness hyperacusis can be modeled and quantified in animals. The results of reaction time and conditioned aversion assays have been interpreted as aversion to sound, but the conclusion that this is due to pain is unclear. Therefore, the subset of hyperacusis known as pain hyperacusis currently has no specific animal model. A tool to measure behavioral changes associated with pain caused by sound, as presented here, is needed to test whether mice can effectively model the subclasses of pain hyperacusis observed in patients.

### Future applications and modifications of detecting pain behavior with composite Facial Grimace and Body Position Scores

The composite Facial Grimace and Body Position Scores presented here provide a valuable initial framework for quantifying pain-related behavioral changes using an unweighted summation of metrics. Notably, certain parameters, such as Ear Ratio and Ear Tip Tilt, exhibited minimal changes in magnitude but significant reductions in variation during painful stimuli, highlighting their potential relevance. Future studies should evaluate the relative contributions of individual parameters in specific pain contexts, allowing for refinement through inclusion, exclusion, or weighting of metrics to optimize the scoring system. It is important to note that the scoring method is designed to provide an inventory that is designed to accommodate further expansion, including new measurements and behavioral categories, such as complex behaviors (e.g., grooming, rearing, exploring, and respiratory rate), which are associated with persistent painful states. (29, 71). Changes in complex behaviors were qualitatively observed in recordings of CGRP-induced migraine and sound exposure but not quantified here. Incorporating these behaviors into a potential third score may enhance the sensitivity and robustness of the method while keeping the simplicity of one camera angle for ease of implementation. By refining the inventory and selecting the most indicative metrics, this approach has the potential to generate a graded pain scale, as demonstrated here by its ability to distinguish between two levels of migraine pain in this initial application.

## Methods

### Animals

Six-week-old C57BL6/J mice of both sexes were used. Both sexes were tested separately. At the time of testing, animals were group-housed in a low noise vivarium. The housing room was maintained at 12:12 h light: dark cycle (7:00 to 19:00). Up to five mice were housed per one filter top shoebox cage (30 × 19 × 13 cm^3^). All procedures were approved by the Johns Hopkins University Animal Care and Use Committee and follow the National Institutes of Health ARRIVE Guidelines.

### Behavioral testing setup design

Individual polycarbonate containers (testing chamber; interior dimensions: 9 x 5x 5 cm) were fabricated onsite and placed in an acoustic chamber prior to testing. The acoustic chamber was lined with LED lights for filming. At all times, the mice were awake and freely moving. Mice were oriented to maintain a profile position via scent cues from air holes positioned at one end of their polycarbonate containers. When multiple mice were recorded simultaneously, two mice of the same sex were placed individually into two chambers oriented to face opposite each other while maintaining a profile position via scent cues from air holes positioned at opposite ends of each container. The mice were prevented from seeing each other by adding a white opaque barrier between the containers. A GoPro Hero 11 Black^TM^ (San Mateo, CA) was centered to capture mice during the protocol. Overhead speakers were fitted into the sound chamber to allow for sound exposure.

### Video Editing

Sound exposure videos were approximately 26 minutes in length and were filmed in 4K resolution at 30 frames per second. Videos were cropped to approximately 1000 x 1500 pixels (height vs width) to analyze one of two mice at a time when multiple mice were recorded simultaneously. The recordings from the CGRP- and saline-injected mice yielded one 10-minute and one 20-minute video filmed in 4K resolution at 30 frames per second. Each video was then cropped using iMovie to focus on a single mouse at a time for the most accurate labeling for each condition. Time segments (epochs) of video were cropped to the desired length to cover a given condition (e.g., CGRP peak epoch, 110 dB SPL, baseline, etc.)

### Pose Estimation using DeepLabCut

The deep neural network was trained to place thirteen markers on the face in each video frame where the face is in profile using DeepLabCut (DLC; (34, 35)). The thirteen markers were labeled inner eye, outer eye, eye up, eye down, nose tip, nose bottom, mouth, ear canal, ear tip, ear center, ear medial, ear lateral, and neck (Figure 1; (23)). DLC was installed on a Windows OS computer with an Intel® Core™ i5-7600K CPU @ 3.80 GHz processor and an Nvidia GeForce GTX 1070 graphics card. ResNet-50-based neural network with default parameters was run for up to 800,000 iterations of training on 816 annotated frames from all cohorts of animals. The dataset was split as 95% for training and 5% for testing. After training, annotated images were visually inspected to ensure the markers were placed on the appropriate key points of the body and face. Data for new videos analyzed were filtered based off a greater than 0.95 likelihood (p-cutoff >= 0.95). A lower limit of at least 30 analyzed frames per video analyzed was used to consider that epoch valid for statistical comparisons, and all recorded time periods fit this criterion.

### Defining Face and Position Parameters

A representative annotated frame showing DLC-generated pose estimated points is shown in Figure 1. Lines between points (skeletons) were created automatically in DLC (Figure 1A). The skeletons used were nose tip to nose bottom, mouth to inner eye, nose bottom to inner eye, inner eye to outer eye, eye up to eye down, inner eye to ear canal, ear canal to ear tip, and medial ear to lateral ear. These skeletons are used to define eye ratio, ear ratio, ear angle, ear position, snout position, and mouth position. Representative images show changes in these parameters between control and pain conditions (Figure 1B). Angles between certain skeletons and ratios of certain skeletons were used to generate measurements of facial parameters, as follows: Ear Ratio was calculated as the ratio of the length of the lines in pixels from Ear Med to Ear Lat and Ear Canal to Ear Tip indicated by white lines (Figure 1C). Ear Position was calculated as the obtuse angle formed by lines connecting Inner Eye, Ear Canal, and Ear Tip with Ear Canal as the vertex as indicated in white (Figure 1D). Ear Tip Tilt was calculated as the angle formed by lines connecting Ear Canal to Ear Center to Ear Tip with Ear Center as the vertex as indicated in white (Figure 1E). Eye Ratio was calculated as the ratio of the length of the lines in pixels from Eye Up to Eye Down and Inner Eye to Outer Eye indicated by white lines (Figure 1F). Snout Position was calculated as the acute angle formed by lines connecting Nose Tip, Inner Eye, and Nose Bottom with Inner Eye as the vertex as indicated in white (Figure 1G). Mouth Position was calculated as the acute angle formed by lines connecting Nose Bottom, Inner Eye, and Mouth with Inner Eye as the vertex as indicated in white (Figure 1H).

The Relative Nose Tip Position was the distance in pixels that the nose tip point was below the (0,0) coordinate in the top left most corner of the chamber, with increasingly negative values representing the mouse’s nose tip at a low place in the chamber (Figure 3A-B). The nose tip pose estimated point was further used to calculate the percentage of analyzed frames where this point is above a horizontal line drawn at −450 pixels (Figure 3C; corresponding to the back of the mouse when all four paws are planted on the chamber floor, ‘Percentage of Nose Tip in Top 1/3 of Chamber’), and to count the incidences when the nose tip crosses a vertical line drawn at 900 pixels (Figure 3D; deliberately set off-center to ensure full turns of the mouse were required to count as an incidence of crossing the vertical line, ‘Vertical Line Crosses of Nose Tip’). These measurements were calculated as the percentage of frames above horizontal line and total number of vertical line crosses, respectively. The horizontal line in Percentage of Nose Tip in Top 1/3 of Chamber was set at −450 because the y-axis pixel count varied from ∼900-1000 based on cropping of the video. The walls of the chamber (top and bottom, including space outside the chamber walls) accounted for 150-300 pixels of that space leaving the interior of the chamber at an average of 900 pixels. Therefore, 1/3 of the chamber is 300, but 150 was added to account for the top chamber wall thus −450. The Percentage of Nose Tip in Top 1/3 of Chamber provides a gross approximation of decreased rearing, exploration of the top portion of the chamber, and climbing behaviors. While

Vertical Line Crosses of Nose Tip capture how often the mouse turns inside the chamber, which is influenced by exploration behavior and total locomotion. Finally, Face Inclination was calculated as the percentage of analyzed frames where the nose tip was below the neck (Figure 3E). Mice in pain tend to hunch forward causing the nose tip to be below the neck point, therefore this measure of face inclination would increase the more the mouse was hunched forward.

### CGRP or Saline Injections

Mice were placed individually into the testing chamber and allowed ten minutes of acclimation to the container. Ten minutes of video was subsequently recorded to measure baseline behavior for each mouse. Mice were then removed from the container and injected with either calcitonin gene-related peptide (CGRP; 100 µg/mL in saline, i.p.) or saline (vehicle) and immediately returned to the container. Since this dose of CGRP induces a migraine that peaks 30 minutes after injection (24), video was recorded from 20 – 40 minutes post injection. In total, 30 minutes of video were recorded for each mouse and was subsequently segmented for analysis as described below.

### Sound Exposure Protocol

Mice received an acclimation period of 30 minutes to their containers the day before the sound exposure protocol. Calibrations within the acoustic chamber were completed each day before noise exposure took place to ensure exposures were accurate and consistent. The two readings checked were 90dB SPL and 120dB SPL. The calibration consisted of taking readings from the corner of each container and the center of the acoustic chamber using a sound level meter model LxT2 (Larson Davis, Inc., Provo, UT). Mice were placed into the same individual polycarbonate containers (described above) within an acoustic chamber fitted with overhead speakers. The mice were unable to look at each other throughout the exposure to eliminate visual cues from the other mouse. The video was centered in the middle capturing both mice during sound exposures. The sound exposure protocol began with ten minutes without sound played from the speakers, then sound was presented in an interleaved protocol ranging from 70 – 120 dB SPL of broadband (2-20 kHz). The sound was played for two minutes followed by two minutes of no sound after each sound presentation. The two-minute period without sound (from 8-10 minutes in each video) was used as the control for each mouse. Sound presentation was interleaved in the following order: 90, 110, 70, 100, 80, 120 dB SPL.

### Data Analysis and Statistics

Filtered CSV files were output once DeepLabCut finished analyzing each video. These files were converted into xlsx files to be quantified. Pairwise comparisons, such as Baseline vs. All – CGRP, were tested using paired, two-tailed, student t-tests. Statistical differences for groups with repeated measures were assessed using repeated measures ANOVA for balanced groups and repeated measures mixed effects model for subgroups tested in longitudinal designs (e.g., CGRP baseline and subgroups within migraine, and sound exposure epochs) and post-hoc Tukey’s correction for multiple comparisons.

## Acknowledgements

This work was supported by the following funding: a Hearing Health Foundation Emerging Research grant to M.B.W., a Johns Hopkins University internal Blaustein Pain Research Fund Grant to B.J.S. and M.B.W., the John Mitchell Jr. Fund to B.J.S., the Geraldine Dietz Fox Endowed Research Fund to E.G. and a National Institute on Deafness and other Communication Disorders (NIDCD) grant 5R01DC016559 to E.G. The David M. Rubenstein Precision Medicine Center of Excellence in Hearing Fund project 2a to EG; the David M. Rubenstein Professorship to EG. We would also like to thank Kevin Psoter, MPA, PhD for consultations of his biostatistical expertise. We thank Drs. Paul Fuchs and Michael Caterina for comments on an earlier version of the manuscript.

---

**Supplemental Figure 1:**
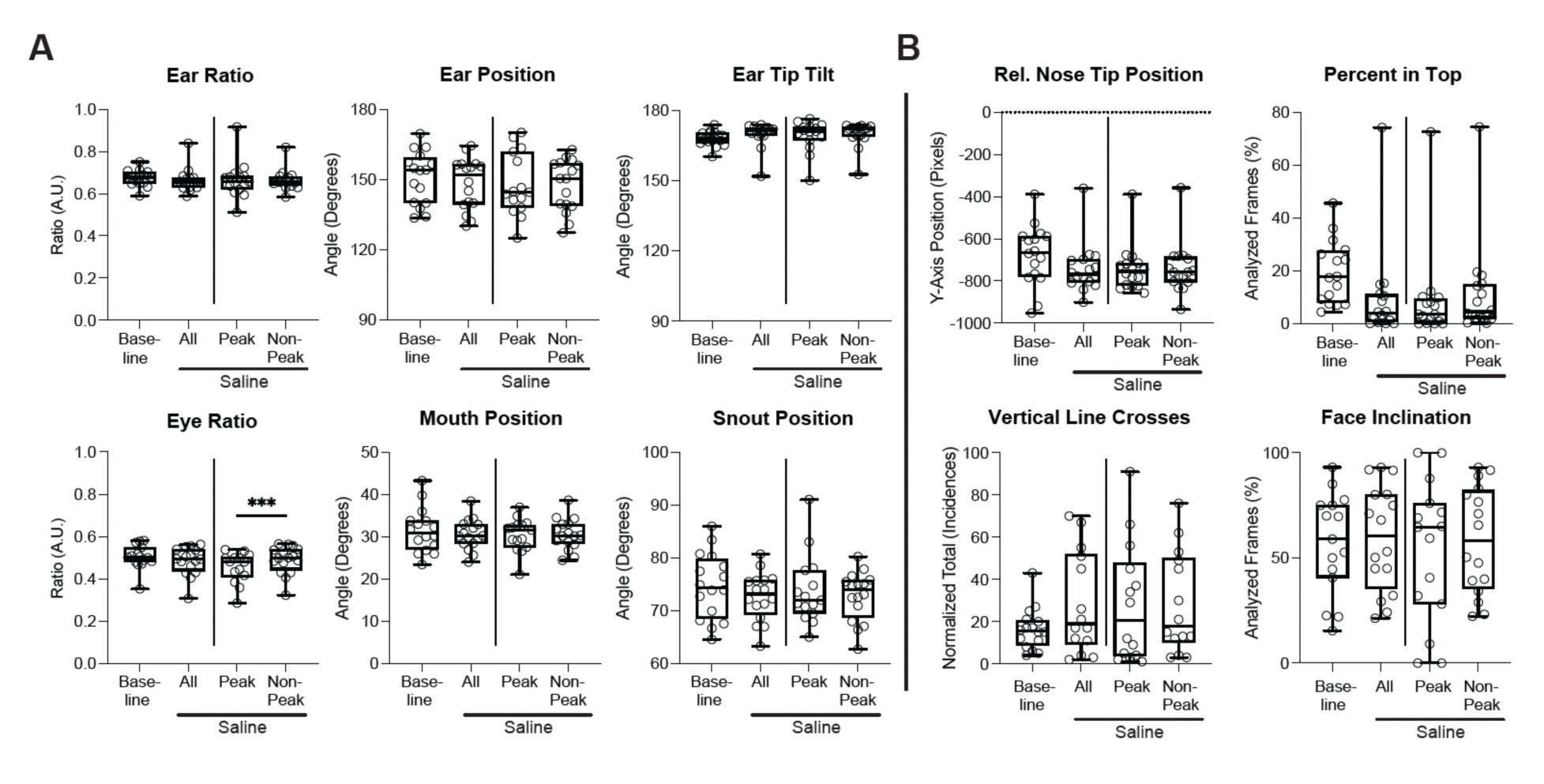
Animals injected with saline do not show changes to facial grimace measurements. **A**) These graphs show the averaged data for each facial parameter indicated. Each dot is the averaged data for the indicated time-period for 1 mouse. Baseline is the averaged data for each mouse for all frames from the Baseline video (10 minutes). ‘*All*’ is the averaged data for each mouse for all frames after CGRP injection. ‘Peak’ is the averaged data from minutes 28-33 and after CGRP injection. ‘*Non-Peak*’ is the averaged data from time outside of the peak time period (minutes 20-26 and minutes 32 to 40). Box and whiskers have a line at the median and the box is the interquartile range with the whiskers extending to the maximum and minimum value of the data. Eye Ratio, repeated measures mixed model, p=0.0465; post hoc comparisons using Tukey’s correction, *Peak* compared to *Non-Peak*, ***, p=0.0003. **B**) These graphs show the averaged data for each body position measurement indicated. Each dot is the averaged data for 1 mouse. *Baseline* is the averaged data for each mouse for all frames from the Baseline video. ‘*All*’ is the averaged data for each mouse for all frames after CGRP injection. ‘*Peak*’ is the averaged data from minutes 27-31 and after CGRP injection. ‘*Non-Peak*’ is the averaged data from time outside of the peak time period (minutes 20-26 and minutes 32 to 40). Box and whiskers have a line at the median and the box is the interquartile range with the whiskers extending to the maximum and minimum value of the data. If no statistics noted, data was not significantly different.

**Supplemental Figure 2:**
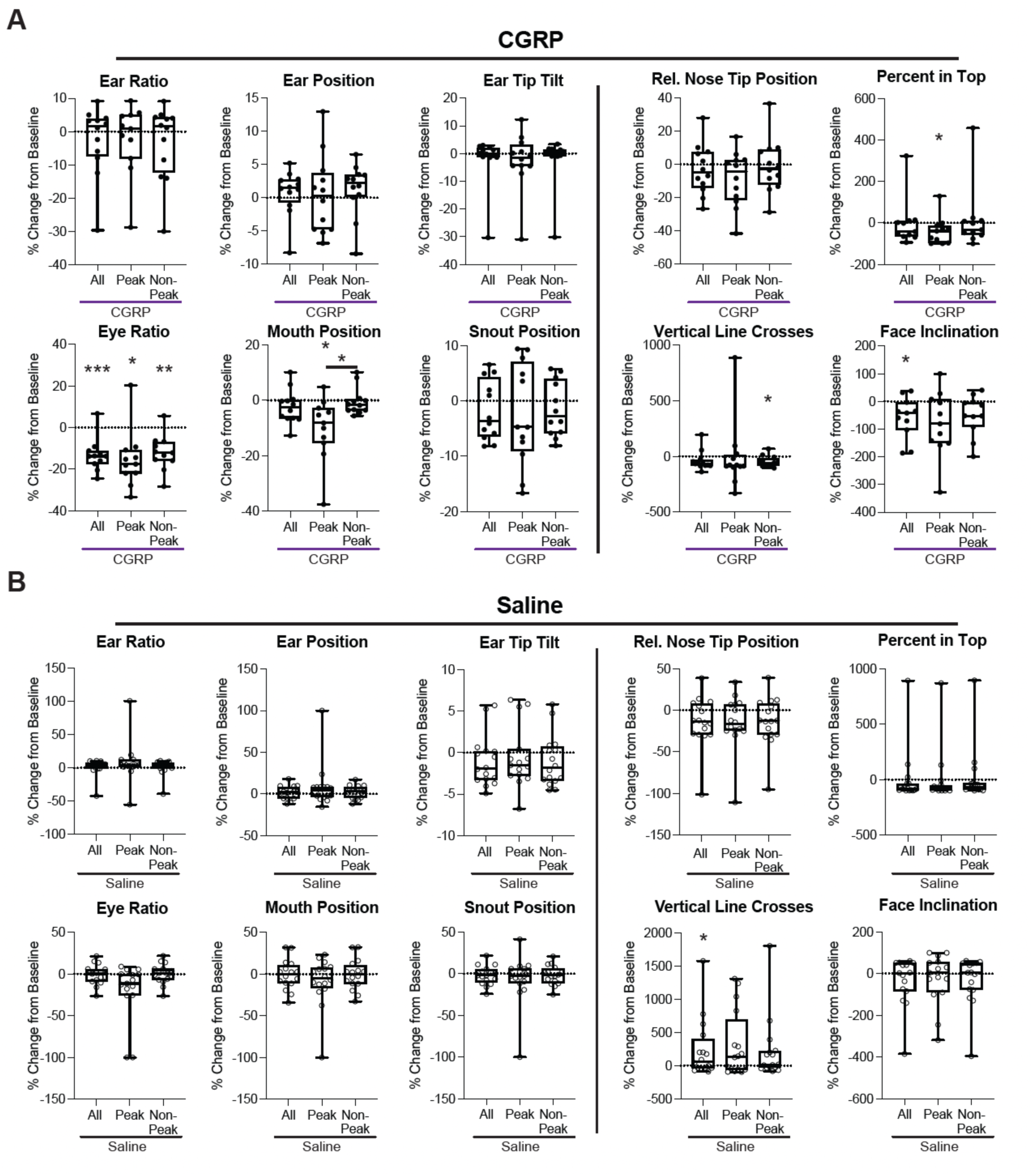
Percentage Change from Baseline for each measurement used to generate Facial Grimace and Body Position Scores for Saline- and CGRP-injected mice. **A**) Graphs of Facial Grimace (left) and Body Position (right) parameters for CGRP-injected mice (N=12). Each dot is the average of one mouse for the indicated time-period after being transformed into a percentage compared to the baseline time-period. If a measurement resulted in a positive change from baseline (Ear Ratio, Ear Position, Rel. Nose Tip Position, Face Inclination), it was inverted so the percentage change from baseline associated with pain was a negative value. Box and whiskers have a line at the median and the box is the interquartile range with the whiskers extending to the maximum and minimum value of the data. Eye Ratio: *Baseline* compared to *All*, paired, two-tailed t-test, p=0.0002; *Baseline* compared to *Peak* and *Non-Peak*, repeated measured mixed model, p= 0.0018, ; post hoc comparisons using Tukey’s correction, *Baseline* compared to *Peak*, *, p=0.0105, *Baseline* compared to *Non-Peak*, **, p=0.0024. Mouth Position: *Baseline* compared to *Peak* and *Non-Peak*, repeated measured mixed model, p= 0.008 ; post hoc comparisons using Tukey’s correction, Baseline compared to *Peak*, *, p=0.0355, *Peak* compared to *Non-Peak*, *, p=0.0417. Percent in Top: *Baseline* compared to *Peak* and *Non-Peak*, repeated measures Friedman Test, p= 0.0435 ; post hoc comparisons using Dunn’s correction, *Baseline* compared to *Peak*, *, p=0.0315. Vertical Line Crosses: *Baseline* compared to *Peak* and *Non-Peak*, repeated measured mixed model, ns, p= 0.6286 ; post hoc comparisons using Tukey’s correction, *Baseline* compared to *Non-Peak*, *, p=0.019. Face Inclination: *Baseline* compared to *All*, paired two-tailed t-test, *, p=0.0255. If no statistics noted, data was not significantly different. **B**) Graphs of Facial Grimace (left) and Body Position (right) parameters for saline-injected mice (N=16). Each open circle is the average of one mouse for the indicated time-period after being transformed into a percentage compared to the baseline time-period. If a measurement resulted in a positive change from baseline (Ear Ratio, Ear Position, Rel. Nose Tip Position, Face Inclination), it was inverted so the percentage change from baseline associated with pain was a negative value. Box and whiskers have a line at the median and the box is the interquartile range with the whiskers extending to the maximum and minimum value of the data. Eye Ratio: *Baseline* compared to *Peak* and *Non-Peak*, repeated measures mixed model, p= 0.0381. Vertical Line Crosses: *Baseline* compared to *All*, Wilcoxon signed-rank test, p=0.0494; *Baseline* compared to *Peak* and *Non-Peak*, repeated measures one-way ANOVA, p=0.0379. If no statistics noted, data was not significantly different. All groups were tested for normality. In cases where the data from all groups was not normally distributed, a Wilcoxon Ranked sum test was used for comparisons of *Baseline* to *All* and a Friedman test was used for the comparison of *Baseline* to *Peak* and *Non-Peak*.

**Supplemental Figure 3:**
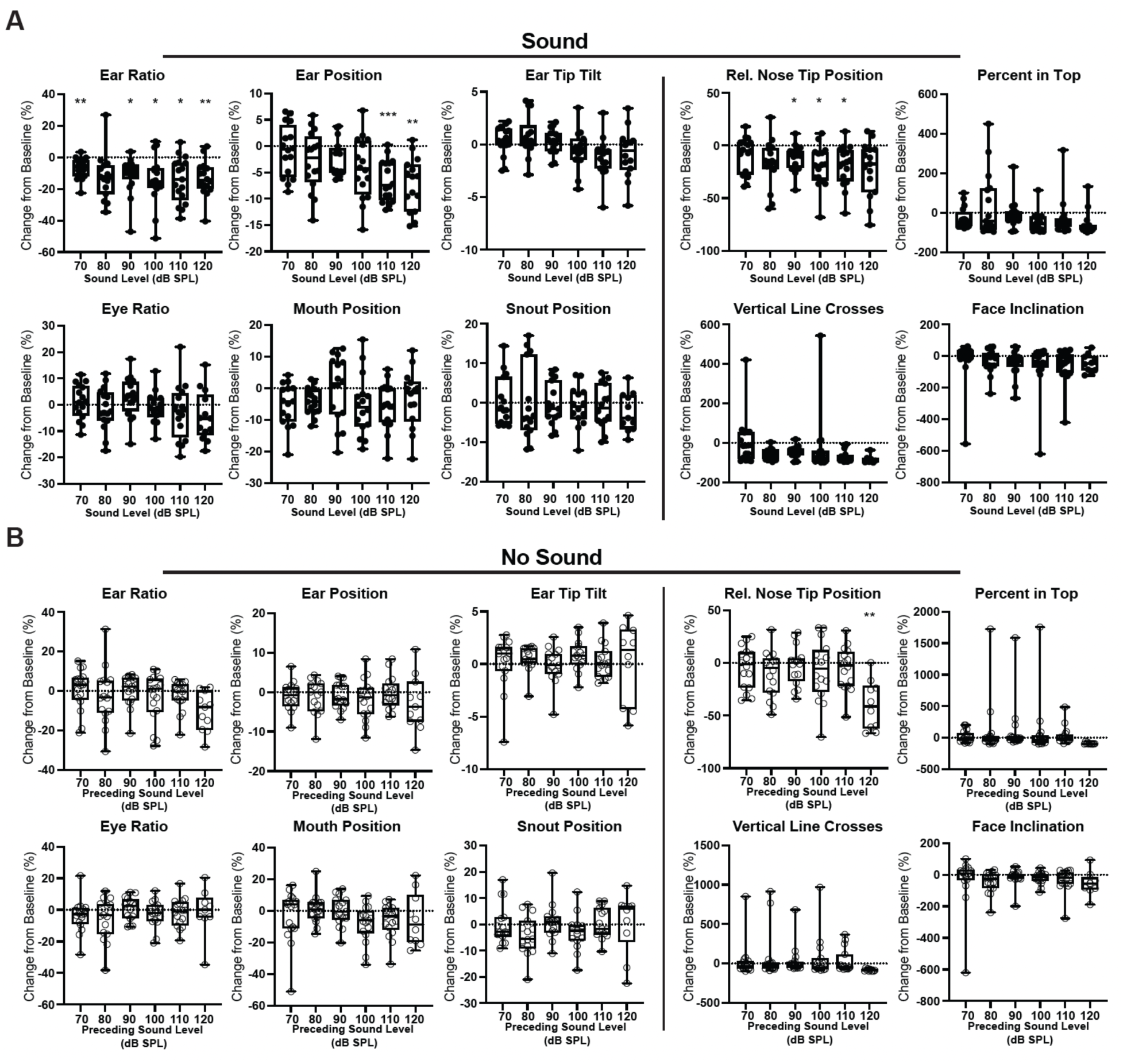
Percentage change from Baseline for each measurement used to generate Facial Grimace and Body Position Scores for Sound- and No-Sound- exposed mice. **A**) Graphs of Facial Grimace (left) and Body Position (right) parameters for sound intervals of the interleaved sound exposure experiment (N=17). Each dot is the average of one mouse for the indicated time-period after being transformed into a percentage compared to the baseline time-period. Sound time-periods have been reordered according to the increasing sound level. If a measurement resulted in a positive change from baseline (Ear Ratio, Ear Position, Rel. Nose Tip Position, Face Inclination), it was inverted so the percentage change from baseline associated with pain was a negative value. Box and whiskers have a line at the median and the box is the interquartile range with the whiskers extending to the maximum and minimum value of the data. Ear Ratio: repeated measures mixed effects model, p=0.0014; post hoc comparisons using Tukey’s correction; baseline compared to 70 dB, **, p=0.0063, baseline compared to 90 dB, *, p=0.0149, baseline compared to 100 dB, *, p=0.0217, baseline compared to 110 dB, *, p=0.0104, baseline compared to 120 dB, **, p=0.0036. Ear Position: repeated measures mixed effects model, p=<0.0001; baseline compared to 110 dB, ***, p=0.0001, baseline compared to 120 dB, **, p=0.001. Ear Tip Tilt: repeated measures mixed effects model, p=0.0139. Eye Ratio: repeated measures mixed effects model, p=0.0469. Rel. Nose Tip Position: repeated measures mixed effects model, p=0.002; post hoc comparisons using Tukey’s correction; baseline compared to 90 dB, *, p=0.0229, baseline compared to 100 dB, *, p=0.0132, baseline compared to 110 dB, *, p=0.0172. Mouth Position, Snout Position, Percent in Top, Vertical Line Crosses, and Face Inclination showed no significant differences between sound levels or from baseline. **B**) Graphs of Facial Grimace (left) and Body Position (right) parameters for no sound intervals of the interleaved sound exposure experiment (N=17). Each dot is the average of one mouse for the indicated time-period after being transformed into a percentage compared to the baseline time-period. No sound time-periods have been reordered according to the preceding sound level. If a measurement resulted in a positive change from baseline (Ear Ratio, Ear Position, Rel. Nose Tip Position, Face Inclination), it was inverted so the percentage change from baseline associated with pain was a negative value. Box and whiskers have a line at the median and the box is the interquartile range with the whiskers extending to the maximum and minimum value of the data. Ear Ratio: repeated measures mixed effects model, p=0.0072; Rel. Nose Tip Position: repeated measures mixed effects model, p=0.0003, post hoc comparisons using Tukey’s correction; baseline compared to post 120 dB, **, p=0.005. Ear Position, Ear Tip Tilt, Eye Ratio, Mouth Position, Snout position, Percent in Top, Vertical line Crosses, and Face Inclination showed no significant differences between no sound periods or from baseline.

**Supplemental Figure 4:**
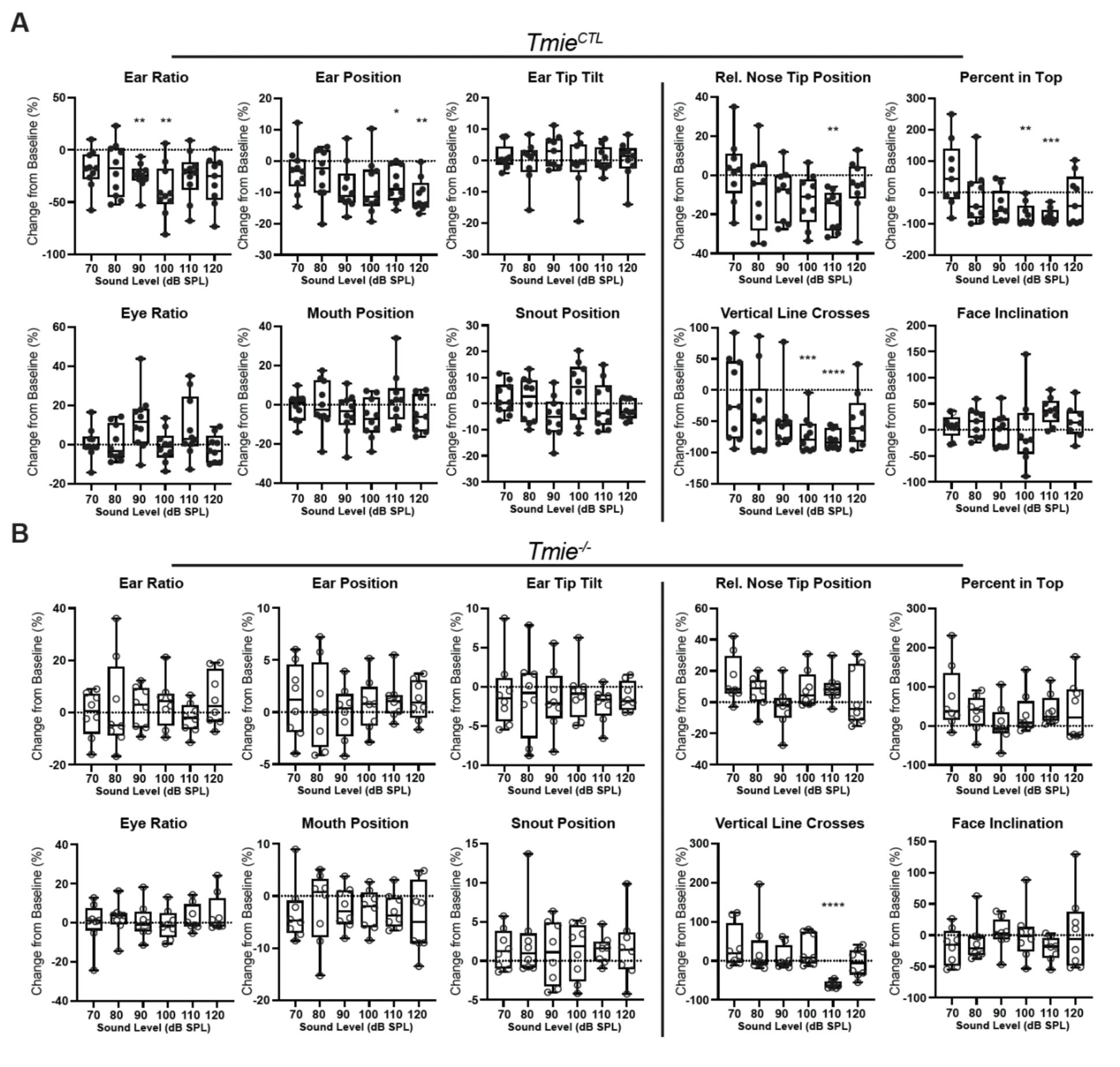
Percentage change from Baseline for each measurement used to generate Facial Grimace and Body Position Scores for *Tmie^CTL^* and *Tmie^-/-^* sound-exposed mice. **A**) Graphs of Facial Grimace (left) and Body Position (right) parameters for sound intervals of *Tmie^CTL^* mice in the interleaved sound exposure experiment (N=10). Each dot is the average of one mouse for the indicated time-period after being transformed into a percentage compared to the baseline time-period. Sound time-periods have been reordered according to increasing sound level. If a measurement resulted in a positive change from baseline (Ear Ratio, Ear Position, Rel. Nose Tip Position, Face Inclination), it was inverted so the percentage change from baseline associated with pain was a negative value. Box and whiskers have a line at the median and the box is the interquartile range with the whiskers extending to the maximum and minimum value of the data. Ear Ratio, *Tmie^CTL^*: repeated measures mixed effects model, p=0.0025; post hoc comparisons using Tukey’s correction, baseline compared to 90 dB, **, p=0.0017, baseline compared to 100 dB, **, p=0.0092. Ear Position, *Tmie^CTL^*: repeated measures mixed effects model, p=0.0349; post hoc comparisons using Tukey’s correction, baseline compared to 110 dB, *, p=0.0237, baseline compared to 120 dB, **, p=0.0035. Eye Ratio, *Tmie^CTL^*: repeated measures mixed effects model, p=0.0321. Rel. Nose Tip Position, *Tmie^CTL^*: repeated measures mixed effects model, p=0.0192; post hoc comparisons using Tukey’s correction, baseline compared to 110 dB, **, p=0.0065. Percent in Top, *Tmie^CTL^*: repeated measures mixed effects model, p=0.0064; post hoc comparisons using Tukey’s correction, baseline compared to 100 dB, **, p=0.0035, baseline compared to 110 dB, ***, p=0.0001. Vertical line crosses, *Tmie^CTL^*: repeated measures mixed effects model, p=0.0062; post hoc comparisons using Tukey’s correction, baseline compared to 100 dB, ***, p=0.0004, baseline compared to 110 dB, ****, p=<0.0001. Face Inclination **B**) Graphs of Facial Grimace (left) and Body Position (right) parameters for sound intervals of *Tmie^-/-^* mice in the interleaved sound exposure experiment (N=8). Each dot is the average of one mouse for the indicated time-period after being transformed into a percentage compared to the baseline time-period. Sound time-periods have been reordered according to the increasing sound level. If a measurement resulted in a positive change from baseline (Ear Ratio, Ear Position, Rel. Nose Tip Position, Face Inclination), it was inverted so the percentage change from baseline associated with pain was a negative value. Box and whiskers have a line at the median and the box is the interquartile range with the whiskers extending to the maximum and minimum value of the data. Rel. Nose Tip Position *Tmie^-/-^*: repeated measures mixed effects model, p=0.0293. Vertical Line Crosses, *Tmie^-/-^*: repeated measures mixed effects model, p=0.0135; post hoc comparisons using Tukey’s correction, baseline compared to 110 dB, ****, p=<0.0001.

## Video Figure Legends

**<u>Supplemental Video 1:</u>** CGRP Experiment Example

10 second video clips from the Baseline recording and 3 segments during the CGRP recording - First 2 Minutes, Peak and Last 2 Minutes. The labels were generated by Deeplabcut and any marker above 0.6 likelihood is shown for visualization. Only markers with a likelihood above 0.95 were used for analysis.

**<u>Supplemental Video 2:</u>** Interleaved Sound Experiment Example

10 second video clips from the original recording of the Baseline, 110 dB SPL, post 110 dB SPL, and 70 dB SPL of a mouse undergoing the interleaved sound protocol. Sound is included in the video.

**<u>Supplemental Video 3:</u>** Labeled, Interleaved Sound Experiment Example

10 second video clips from the labeled recording of the Baseline, 110 dB SPL, post 110 dB SPL, and 70 dB SPL time periods of a mouse undergoing the interleaved sound protocol. The labeled video is from the same time periods from the same mouse shown in Supplemental Video 2. The labels were generated by Deeplabcut and any marker above 0.6 likelihood is shown for visualization. Only markers with a likelihood above 0.95 were used for analysis.

**<u>Supplemental Video 4:</u>** *Tmie^CTL^* Example Video

10 second video clips from an original recording of the Baseline and 110 dB SPL time periods of a *Tmie^CTL^* mouse undergoing the interleaved sound protocol with sound included. The original clips are followed by the corresponding labeled video. The labels were generated by Deeplabcut and any marker above 0.6 likelihood is shown for visualization. Only markers with a likelihood above 0.95 were used for analysis.

**<u>Supplemental Video 5:</u>** *Tmie^-/-^* Example Video

10 second video clips from an original recording of the Baseline and 110 dB SPL time periods of a *Tmie^-/-^* mouse undergoing the interleaved sound protocol with sound included. The original clips are followed by the corresponding labeled video. The labels were generated by Deeplabcut and any marker above 0.6 likelihood is shown for visualization. Only markers with a likelihood above 0.95 were used for analysis.

## References

1. H. E. von Gierke, D. R. Brown, Wright-Patterson AFB, Ohio. Benox Report; an Exploratory Study of the Biological Effects of Noise 47 (1953).

2. G. Frank, M. Kössl, The acoustic two-tone distortions 2f1-f2 and f2-f1 and their possible relation to changes in the operating point of the cochlear amplifier. Hear Res 98, 104–115 (1996).

3. L. P. Sherlock, C. Formby, Estimates of loudness, loudness discomfort, and the auditory dynamic range: normative estimates, comparison of procedures, and test-retest reliability. J Am Acad Audiol 16, 85–100 (2005).

4. S. Manohar, H. J. Adler, K. Radziwon, R. Salvi, Interaction of auditory and pain pathways: Effects of stimulus intensity, hearing loss and opioid signaling. Hear Res 393, 108012 (2020).

5. R. S. Tyler, et al., A review of hyperacusis and future directions: part I. Definitions and manifestations. Am J Audiol 23, 402–419 (2014).

6. M. Pienkowski, et al., A review of hyperacusis and future directions: part II. Measurement, mechanisms, and treatment. Am J Audiol 23, 420–436 (2014).

7. B. Pollard, Clinical advancements for managing hyperacusis with pain. The Hearing Journal 72, 10–12 (2019).

8. R. Salvi, G.-D. Chen, S. Manohar, Hyperacusis: Loudness intolerance, fear, annoyance and pain. Hear Res 426, 108648 (2022).

9. K. N. Jahn, S. T. Kashiwagura, M. S. Yousuf, Clinical phenotype and management of sound-induced pain: Insights from adults with pain hyperacusis. J Pain 27, 104741 (2025).

10. J. J. Song, et al., Hyperacusis-associated pathological resting-state brain oscillations in the tinnitus brain: a hyperresponsiveness network with paradoxically inactive auditory cortex. Brain Struct Funct 219, 1113–28 (2014).

11. W. Zhou, et al., Sound induces analgesia through corticothalamic circuits. Science 377, 198–204 (2022).

12. K. K. Clayton, et al., Sound elicits stereotyped facial movements that provide a sensitive index of hearing abilities in mice. Curr Biol 34, 1605–1620.e5 (2024).

13. K. Radziwon, R. Salvi, Using auditory reaction time to measure loudness growth in rats. Hear Res 395, 108026 (2020).

14. S. S. Smith, et al., Objective autonomic signatures of tinnitus and sound sensitivity disorders. Sci Transl Med 17, eadp1934. (2025).

15. A. E. Hickox, M. C. Liberman, Is noise-induced cochlear neuropathy key to the generation of hyperacusis or tinnitus? J Neurophysiol 111, 552–564 (2014).

16. R. H. Salloum, et al., Untangling the effects of tinnitus and hypersensitivity to sound (hyperacusis) in the gap detection test. Hear Res 331, 92–100 (2016).

17. J. R. Deuis, L. S. Dvorakova, I. Vetter, Methods Used to Evaluate Pain Behaviors in Rodents. Front Mol Neurosci 10, 284 (2017).

18. K. J. Sufka, Conditioned place preference paradigm: a novel approach for analgesic drug assessment against chronic pain. Pain 58, 355–366 (1994).

19. Z. Zhang, et al., Automated preclinical detection of mechanical pain hypersensitivity and analgesia. Pain 163, 2326–2336 (2022).

20. M. Bohic, et al., Mapping the neuroethological signatures of pain, analgesia, and recovery in mice. Neuron 111, 2811–2830.e8 (2023).

21. D. J. Langford, et al., Coding of facial expressions of pain in the laboratory mouse. Nat Methods 7, 447–449 (2010).

22. E. S. McCoy, et al., Development of PainFace software to simplify, standardize, and scale up mouse grimace analyses. Pain 165, 1793–1805 (2024).

23. O. Le Moëne, M. Larsson, A New Tool for Quantifying Mouse Facial Expressions. eNeuro 10, ENEURO.0349-22.2022 (2023).

24. B. J. Rea, et al., Peripherally administered calcitonin gene-related peptide induces spontaneous pain in mice: implications for migraine. Pain 159, 2306–2317 (2018).

25. A. H. Tuttle, et al., A deep neural network to assess spontaneous pain from mouse facial expressions. Mol Pain 14, 1744806918763658 (2018).

26. C.-Y. Chiang, et al., Deep Learning-Based Grimace Scoring Is Comparable to Human Scoring in a Mouse Migraine Model. J Pers Med 12 (2022).

27. E. J. Cobos, et al., Inflammation-induced decrease in voluntary wheel running in mice: a nonreflexive test for evaluating inflammatory pain and analgesia. Pain 153, 876–884 (2012).

28. G. J. Bennett, What is spontaneous pain and who has it? J Pain 13, 921–929 (2012).

29. P. V. Turner, D. S. Pang, J. L. Lofgren, A Review of Pain Assessment Methods in Laboratory Rodents. Comp Med 69, 451–467 (2019).

30. M. R. Gleason, et al., The transmembrane inner ear (Tmie) protein is essential for normal hearing and balance in the zebrafish. Proc Natl Acad Sci U S A 106, 21347–21352 (2009).

31. Z. Wu, et al., Mechanosensory hair cells express two molecularly distinct mechanotransduction channels. Nat Neurosci 20, 24–33 (2017).

32. J. R. Holt, et al., Putting the Pieces Together: the Hair Cell Transduction Complex. J Assoc Res Otolaryngol 22, 601–608 (2021).

33. R. Fettiplace, D. N. Furness, M. Beurg, The conductance and organization of the TMC1-containing mechanotransducer channel complex in auditory hair cells. Proc Natl Acad Sci U S A 119, e2210849119 (2022).

34. A. Mathis, et al., DeepLabCut: markerless pose estimation of user-defined body parts with deep learning. Nat Neurosci 21, 1281–1289 (2018).

35. T. Nath, et al., Using DeepLabCut for 3D markerless pose estimation across species and behaviors. Nat Protoc 14, 2152–2176 (2019).

36. S. M. Stubsjøen, et al., Exploring non-invasive methods to assess pain in sheep. Physiol Behav 98, 640–648 (2009).

37. J. Stracke, B. Bert, H. Fink, J. Böhner, Assessment of stress in laboratory beagle dogs constrained by a Pavlov sling. ALTEX 28, 317–325 (2011).

38. A. Schroeer, F. I. Corona-Strauss, R. Hannemann, S. A. Hackley, D. J. Strauss, The vestigial pinna-orienting system in humans briefly suppresses superior auricular muscle activity during reflexive orienting toward auditory stimuli. J Neurophysiol 132, 514–526 (2024).

39. E. Friauf, H. Herbert, Topographic organization of facial motoneurons to individual pinna muscles in rat (Rattus rattus) and bat (Rousettus aegyptiacus). J Comp Neurol 240, 161–170 (1985).

40. J. E. Heuser, T. I. Tenkova, Introducing a mammalian nerve-muscle preparation ideal for physiology and microscopy, the transverse auricular muscle in the ear of the mouse. Neuroscience 439, 80–105 (2020).

41. L. H. Lassen, et al., CGRP may play a causative role in migraine. Cephalalgia 22, 54–61 (2002).

42. B. N. Mason, et al., Induction of Migraine-Like Photophobic Behavior in Mice by Both Peripheral and Central CGRP Mechanisms. J Neurosci 37, 204–216 (2017).

43. Hasriadi, P. W. D. Wasana, O. Vajragupta, P. Rojsitthisak, P. Towiwat, Automated home-cage for the evaluation of innate non-reflexive pain behaviors in a mouse model of inflammatory pain. Sci Rep 11, 12240 (2021).

44. M. J. O. Boedts, Tympanic Resonance Hypothesis. Front Neurol 11, 14 (2020).

45. R. T. Bottom, Y. Xu, C. Siebald, J. Jung, U. Müller, Defects in hair cells disrupt the development of auditory peripheral circuitry. Nat Commun 15, 10899 (2024).

46. P. Fournier, et al., Contraction of the stapedius and tensor tympani muscles explored by tympanometry and pressure measurement in the external auditory canal. Hear Res 420, 108509 (2022).

47. V. Rinaldi, M. Cappadona, M. Gaffuri, S. Torretta, L. Pignataro, Chorda tympani nerve, may it have a role in stabilizing middle ear pressure? Medical hypotheses 80, 726–727 (2013).

48. T. J. Rockley, W. M. Hawke, The middle ear as a baroreceptor. Acta Otolaryngol 112, 816–823 (1992).

49. M. Westcott, et al., Tonic tensor tympani syndrome in tinnitus and hyperacusis patients: a multi-clinic prevalence study. Noise Health 15, 117–128 (2013).

50. A. E. Sutton, R. De Jong, G. Kwartowitz, “Tensor Tympani Syndrome.” in StatPearls, (StatPearls Publishing, 2025).

51. A. S. Filippidis, S. G. Zarogiannis, A. Randich, T. J. Ness, S. Matalon, Assessment of locomotion in chlorine exposed mice by computer vision and neural networks. J Appl Physiol (1985) 112, 1064–1072 (2012).

52. A. M. Kolstad, R. M. Rodriguiz, C. J. Kim, L. P. Hale, Effect of pain management on immunization efficacy in mice. J Am Assoc Lab Anim Sci 51, 448–457 (2012).

53. D. Vuralli, A.-S. Wattiez, A. F. Russo, H. Bolay, Behavioral and cognitive animal models in headache research. J Headache Pain 20, 11 (2019).

54. A. Ashiquzzaman, et al., MoSeq based 3D behavioral profiling uncovers neuropathic behavior changes in diabetic mouse model. Sci Rep 15, 15114 (2025).

55. C. Weinreb, et al., Keypoint-MoSeq: parsing behavior by linking point tracking to pose dynamics. Nat Methods 21, 1329–1339 (2024).

56. W. A. Yost, M. C. Killion, Hearing thresholds. Encyclopedia of acoustics 3, 1545–1554 (1997).

57. S. E. Shore, C. Wu, Mechanisms of Noise-Induced Tinnitus: Insights from Cellular Studies. Neuron 103, 8–20 (2019).

58. S. Mukerji, A. M. Windsor, D. J. Lee, Auditory brainstem circuits that mediate the middle ear muscle reflex. Trends Amplif 14, 170–191 (2010).

59. C. Liu, E. Glowatzki, P. A. Fuchs, Unmyelinated type II afferent neurons report cochlear damage. Proc Natl Acad Sci U S A 112, 12723–7 (2015).

60. E. N. Flores, et al., A non-canonical pathway from the cochlea to brain signals tissue-damaging noise. Curr Biol 25, 606–12 (2015).

61. N. Nowak, M. B. Wood, E. Glowatzki, P. A. Fuchs, Prior Acoustic Trauma Alters Type II Afferent Activity in the Mouse Cochlea. eNeuro 8, ENEURO.0383-21.2021 (2021).

62. T. Nagai, T. Tono, Encapsulated nerve corpuscles in the human tympanic membrane. Arch Otorhinolaryngol 246, 169–172 (1989).

63. K. Limakatso, G. J. Bedwell, V. J. Madden, R. Parker, The prevalence and risk factors for phantom limb pain in people with amputations: A systematic review and meta-analysis. PLoS One 15, e0240431 (2020).

64. M. McGill, et al., Neural signatures of auditory hypersensitivity following acoustic trauma. Elife 11 (2022).

65. M. Qian, et al., The quantification and mRNA expression levels of cochlear synapses in C57BL/6j mice following repeated exposure to noise. Acta Otolaryngol 144, 558–564 (2024).

66. K. A. Fernandez, et al., Noise-induced Cochlear Synaptopathy with and Without Sensory Cell Loss. Neuroscience 427, 43–57 (2020).

67. M. B. Wood, et al., Acoustic Trauma Increases Ribbon Number and Size in Outer Hair Cells of the Mouse Cochlea. J Assoc Res Otolaryngol 22, 19–31 (2021).

68. K. M. Schrode, M. L. Dent, A. M. Lauer, Sources of variability in auditory brainstem response thresholds in a mouse model of noise-induced hearing loss. J Acoust Soc Am 152, 3576 (2022).

69. S. Manohar, G.-D. Chen, L. Li, X. Liu, R. Salvi, Chronic stress induced loudness hyperacusis, sound avoidance and auditory cortex hyperactivity. Hear Res 431, 108726 (2023).

70. S. Manohar, J. Spoth, K. Radziwon, B. D. Auerbach, R. Salvi, Noise-induced hearing loss induces loudness intolerance in a rat Active Sound Avoidance Paradigm (ASAP). Hear Res 353, 197–203 (2017).

71. N. R. Council, D. on Earth, I. for L. A. Research, C. on Recognition, A. of P. in L. Animals, Recognition and alleviation of pain in laboratory animals. (2010

